# The root knot nematode effector MiEFF186 interacts with a splicing factor to reshape plant transcriptome

**DOI:** 10.64898/2026.10.12.758410

**Authors:** Sarah Ranty-Roby, Karine Mulet, Salomé Soulé, Laurent Camborde, Kaiwei Huang, Piao Lei, Jess Philippon, Claire Caravel, Pauline Foubert, Maria Belen de Luca, Carine Huard-Chauveau, Stéphanie Jaubert, Martin D. Crespi, Jianlong Zhao, Jérémie Bazin, Bruno Favery, Michaël Quentin

## Abstract

The root-knot nematode *Meloidogyne incognita* secretes specific effectors to manipulate root cells and induce their redifferentiation into multinucleate feeding ‘giant cells’, which are essential for its development. A dedicated screen identified the *Meloidogyne incognita* MiEFF186a effector as a regulator of alternative splicing, a crucial mechanism of transcriptional gene regulation whereby a single gene can produce multiple mature mRNA variants, in plants. EFF186 is expressed in the esophageal glands of nematode juveniles, and *in planta*, it localizes to the nucleus and accumulates in the nucleolus. A yeast two-hybrid approach identified the nucleolar G-patch protein GDP1/CIS2 as a MiEFF186a target in tomato and Arabidopsis. Transcriptomic analyses of *Arabidopsis* roots ectopically expressing *MiEFF186a* or impaired in GDP1/CIS2 function revealed changes in both gene expression and alternative splicing, with partially overlapping responses. Finally, alteration of *GDP1/CIS2* expression by virus-induced gene silencing in *Solanaceae* or in *Arabidopsis gdp1* mutants affects susceptibility to *M. incognita*. Together, these results identify GDP1/CIS2 as a host target of MiEFF186a and support a role for these proteins in the modulation of host RNA metabolism during nematode parasitism.

## 1 | INTRODUCTION

In eukaryotes, splicing consists in the removal of non-coding sequences, introns, from the precursor messenger RNA (pre-mRNA) molecule to create a mature mRNA ready for protein translation. It is allowed by the spliceosome, a large and dynamic complex made of RNA and proteins. The spliceosome comprises five uridine (U)-rich small nuclear ribonucleoprotein subcomplexes (snRNPs; U1, U2, U4, U5, and U6), and contains several additional proteins including trans-acting splicing factors (Plaschka *et al*., 2018). The U1, U2, U4 and U5 share a heptameric core consisting of Sm proteins (SmD1, D2, D3, E, F and G), while U6 is made of related Sm-like proteins (LSm2-8). Alternative splicing (AS) is a process by which a single gene’s pre-mRNA transcript is cut and re-joined in different ways, allowing to produce multiple mature mRNA isoforms (Reddy, 2007). AS may generates protein variants, altering domain composition, catalytic residues, or changing protein stability or subcellular localization. Aberrant transcript isoforms may also be produced that trigger nonsense-mediated mRNA decay (NMD), or are retained in the nucleus, permitting a rapid downregulation of active transcript levels (Tognacca *et al*., 2023; Jabre *et al*., 2026). AS is essential for the modulation of cellular activities in response to various physiological and developmental signals. In plants, 40-70% of multi-exonic genes produce more than one transcript by AS, with intron retention being the most frequent event (Chamala *et al*., 2015). AS is controlled by cis-regulatory sequences within the exons or introns (e.g. 5’ and 3’ splice sites, splicing enhancers and silencers) and trans-acting factors including RNA-binding proteins (RBPs, *e.g.* splicing factors) (Reddy, 2007). AS contributes to plant development and to biotic and abiotic stress responses, allowing plants to rapidly adapt to their environment (Hewezi, 2024; Laloum *et al*., 2018). Plant infection is associated with rapid and specific changes in plant AS patterns (Rigo *et al*., 2019; Hewezi, 2024; Godinho *et al*., 2025; Gao *et al*., 2026). Several splicing-regulatory effectors (SREs) secreted from bacteria, oomycetes and nematodes RBPs and splicing factors, in order to modulate plant immunity (Hewezi, 2024; Ranty-Roby *et al*., 2024; Godinho *et al*., 2025; Gao *et al*., 2026).

Root-knot nematodes (RKNs; *Meloidogyne* spp.) are microscopic worms capable of inducing spectacular morphological and physiological changes in the root cells of several thousand host plants (Jones *et al*., 2013). They are obligate biotrophic endoparasites. Within the central cylinder, juveniles select five to seven vascular parenchyma cells and induce their redifferentiation into specialized feeding cells known as “giant cells”. These hypertrophied, multinucleated, and metabolically hyperactive cells serve as a permanent and exclusive source of nutrients for the nematode. These giant feeding cells are therefore essential for nematode development and reproduction (Favery *et al*., 2021; Rutter *et al*., 2022). At the same time, the cells surrounding the giant cells proliferate and both the phloem and xylem expand. This hyperplasia of root tissues leads to the formation of the gall, which is characteristic of *Meloidogyne* infection (Favery *et al*., 2021; Rutter *et al*., 2022). Giant cells and gall formation requires an extensive regulation of gene’s expression, and AS was shown to contribute to modulating host transcriptome (Mejias *et al*., 2021; Mejias *et al*., 2022; Ozdemir *et al*., 2024). Proteinaceous effectors, produced in esophageal glands of the RKN and secreted *in planta* through a syringe-like stylet, are instrumental in hijacking host transcriptional machinery (Bali & Gleason, 2024; Rutter *et al*., 2022). Recently, two RKN SREs were described. A conserved RKN effector, EFF18, contributes to diversifying host cell transcriptome during GC formation. EFF18 was shown to interact with the core spliceosomal protein SmD1, in Arabidopsis, *Nicotinana benthamiana* and tomato and contributes to diversifying the host transcriptome during giant cell formation. Perturbating SmD1 expression decreases plant susceptibility to *M. incognita* and impaires giant cells development (Mejias *et al*., 2021 and 2022). Another RKN effector, MiMSP8, was shown to interact with the tomato SlU2AF35, a subunit of the U2-associated splicing factor U2AF (Chen *et al*., 2025). Silencing SlU2AF35 impacted tomato susceptibility to RKN. MiMSP8 was shown to disturb U2AF assembly and its binding to a subset of pre-mRNAs (Chen *et al*., 2025). Targeting AS thus provides a powerful means for nematodes to alter the plant transcriptome and evade defenses.

In this study, we screened several nuclear effectors from *M. incognita* to assess their role in regulating AS and identified a novel nucleolar SRE, MiEFF186a. We found that MiEFF186a binds to CRY2 INTERACTING SPLICING FACTOR 2 (CIS2), a nucleolar G-patch domain-containing protein also known as G-PATCH DOMAIN PROTEIN 1 (GDP1) and involved in both alternative splicing and pre-rRNA processing in *Arabidopsis* (Kojima *et al*., 2018; Zhao *et al*., 2022). Here, we show that MiEFF186a and GDP1/CIS2 partially colocalize in the nucleolus and that loss of GDP1/CIS2 affects plant susceptibility to *M. incognita*. Transcriptomic analyses further reveal that *MiEFF186a* expression and *GDP1/CIS2* disruption affect both gene expression and alternative splicing, with partially overlapping responses. Therefore, MiEFF186a likely contributes to nematode infection by reprogramming host RNA metabolism through its interaction with GDP1/CIS2.

## 2 | RESULTS

### 2.1 | Identification of splicing regulatory effectors from *M. incognita*

To search for new effectors involved in the regulation of AS during RKN parasitism, we used the luminescence-based AS reporter system in *N. benthamiana* that was previously established to characterize *Phytophthora infestans* SREs (Huang *et al*., 2020). These plants express part of a tomato receptor-like protein kinase (*RLPK*; *Solyc03g007210*) gene with an intron, fused to the luciferase reporter gene (LUC). When the intron is removed through splicing, the transcript produces a functional LUC whose activity can be quantified, while if the intron which has a premature termination codon is retained, no LUC is produced. Among described *M. incognita* candidate effectors, 13 were selected owing a nuclear localization signal (NLS) and/or a nucleolar localization signal (NoLS) that could putatively target the macromolecular splicing machinery (Fig 1a; Fig. S1). *Agrobacterium tumefaciens* strains carrying GFP-effector fusions were infiltrated into *N. benthamiana* p35S::RLPK-LUC leaves. GFP was used as a negative control, while the *P. infestans* SRE3 and SRE4 (Huang *et al*., 2020) and MiEFF18a (Mejias *et al*., 2021) were used as positive controls triggering intron retention. LUC activity was quantified at 48 hours post infiltration. As SRE3, SRE4 and MiEFF18a, only *Minc3s02248g29071* significantly and reproducibly affected AS in the splicing reporter assays (adjusted p-value<0.01; Fig. 1b). RT-qPCRs were then performed to verify that the candidate RKN SRE affected AS and not just the expression level of the RLPK-LUC transcript. MiEFF18 and the GFP were used as a positive and negative control, respectively. RT-qPCR confirmed that Minc3s02248g29071 is a true SRE, while Minc03325 is not (Fig. S2). Minc3s02248g29071, which shows a significant effect on regulating AS was considered for further studies and will later be referred to as MiEFF186a.

**Figure 1.**
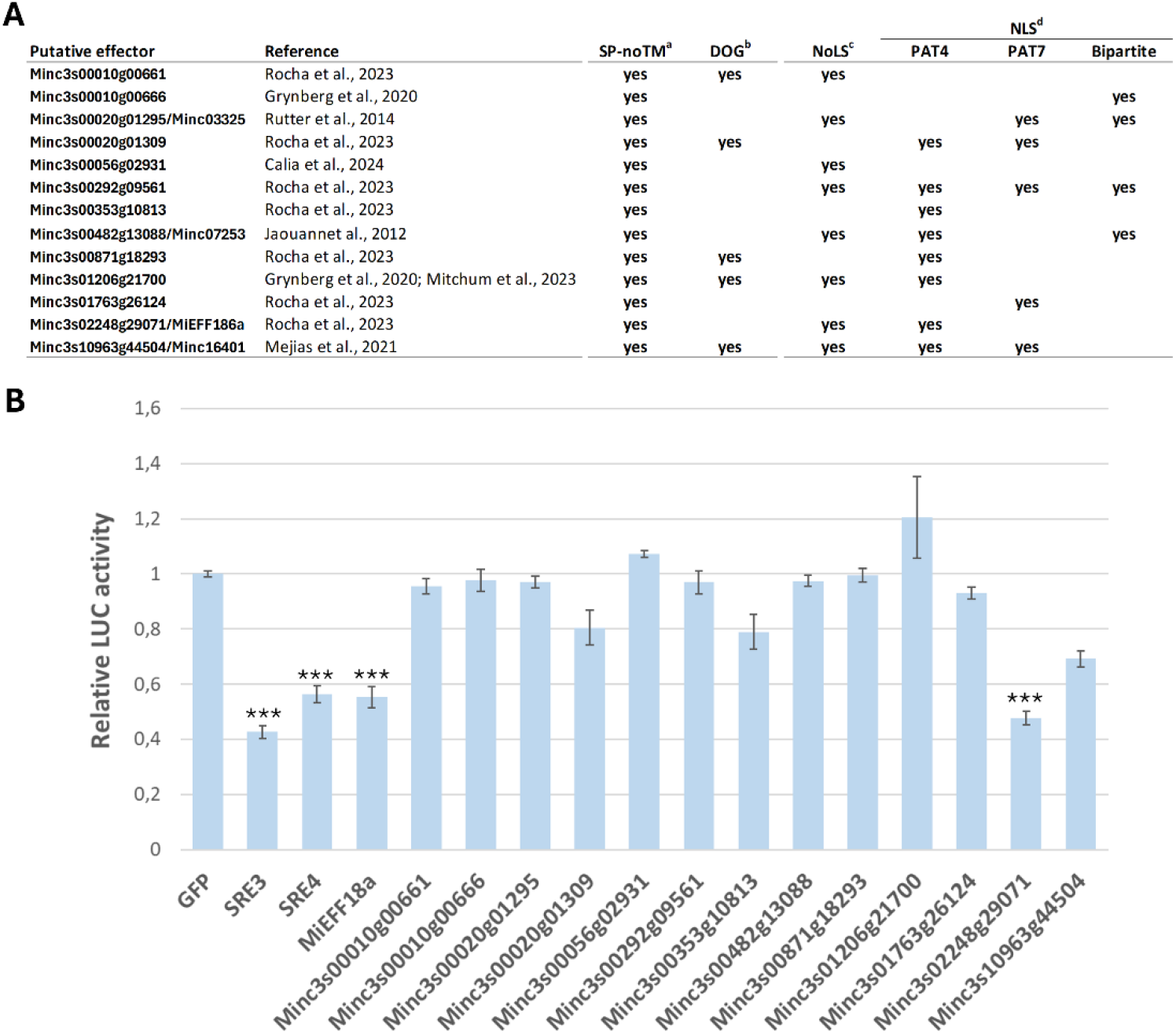
Screening with a splicing reporter assay identified a new *Meloidogyne incognita* splicing regulatory effector (SRE). **A,** Candidate nuclear effectors tested for AS modulation. ^a^ Protein containing a predicted N-terminal signal peptide (SP) and lacking transmembrane domains (noTM) according to the signalP program (Almagro Armenteros *et al*., 2019); ^b^ Candidate effector carrying a Mel-DOG box characteristic of dorsal gland effectors (da Rocha *et al*., 2021); ^c^ Nucleolar Localisation Signals (NoLS) were predicted with NoD (Scott *et al*., 2011); ^d^ Nuclear Localization Signals (NLS) were predicted with PSORT (Nakai & Horton, 1999). **B,** Screen with a reporter assay identified Minc3s02248g29071/MiEFF18a as a new SRE. Leaves of p35S::RLPK-LUC transgenic *N. benthamiana* plants were infiltrated with A. tumefaciens strains containing a GFP-effector or GFP-control construct, and LUC activity was quantified at 48hpi. Leaf discs from 8 individual leaves were collected for LUC activity measurement, and relative LUC activity was normalized to the activity of the corresponding GFP control. Means and standard errors are shown (n=8 to 45). Only SRE, SRE4; MiEFF18 and Minc3s002248g2971 displayed significant SRE activity when compared to GFP (*** adjusted p-value<0.001; Dunn-Bonferroni test).

### 2.2 | MiEFF186 is a conserved RKN effector

The MiEFF186a protein is 235 amino-acids long and possesses a signal peptide (SP), a nucleolar localization signal (NoLS). It shows no similarity to any domain available in public databases (Fig. 2A). It possesses a consensus disorder, lacking a stable three-dimensional structure, which is often found in proteins interacting with different partners (Trivedi & Nagarajaram, 2022). The MiEFF186 sequence was searched for homologs within the *M.* incognita genome, resulting in the identification of six homologs (Fig. 2B; Supplementary Fig. S3). The EFF186 genes are specific to RKNs and, using BLAST tools, could be recovered in the RKN genome of *M. incognita*, *M. javanica*, *M. arenaria*, *M. hapla*, *M. enterolobii*, *M. floridensis*, and *M. luci*, but not in the genomes of other plant parasitic nematodes or other organisms (Fig. 2B; Supplementary Fig. S3 and S4). A phylogenetic tree indicated that all taxa within the tree exhibit a high degree of genetic relatedness (Fig. 2B; Supplementary Fig. S4). Thus EFF186s are conserved and exclusive of RKNs. Noteworthy, MiEFF186a was identified in esophageal gland-enriched samples through both transcriptomic and glycoproteomic analyses (Rocha *et al*., 2023), suggesting that it is secreted *in planta*, and that it is likely involved in parasitism and functions as an effector. Consistently, analysis of available RNAseq data showed MiEFF186a genes is more strongly expressed during early parasitic stages (Fig. 2C; Blanc-Mathieu *et al*., 2017).

**Figure 2.**
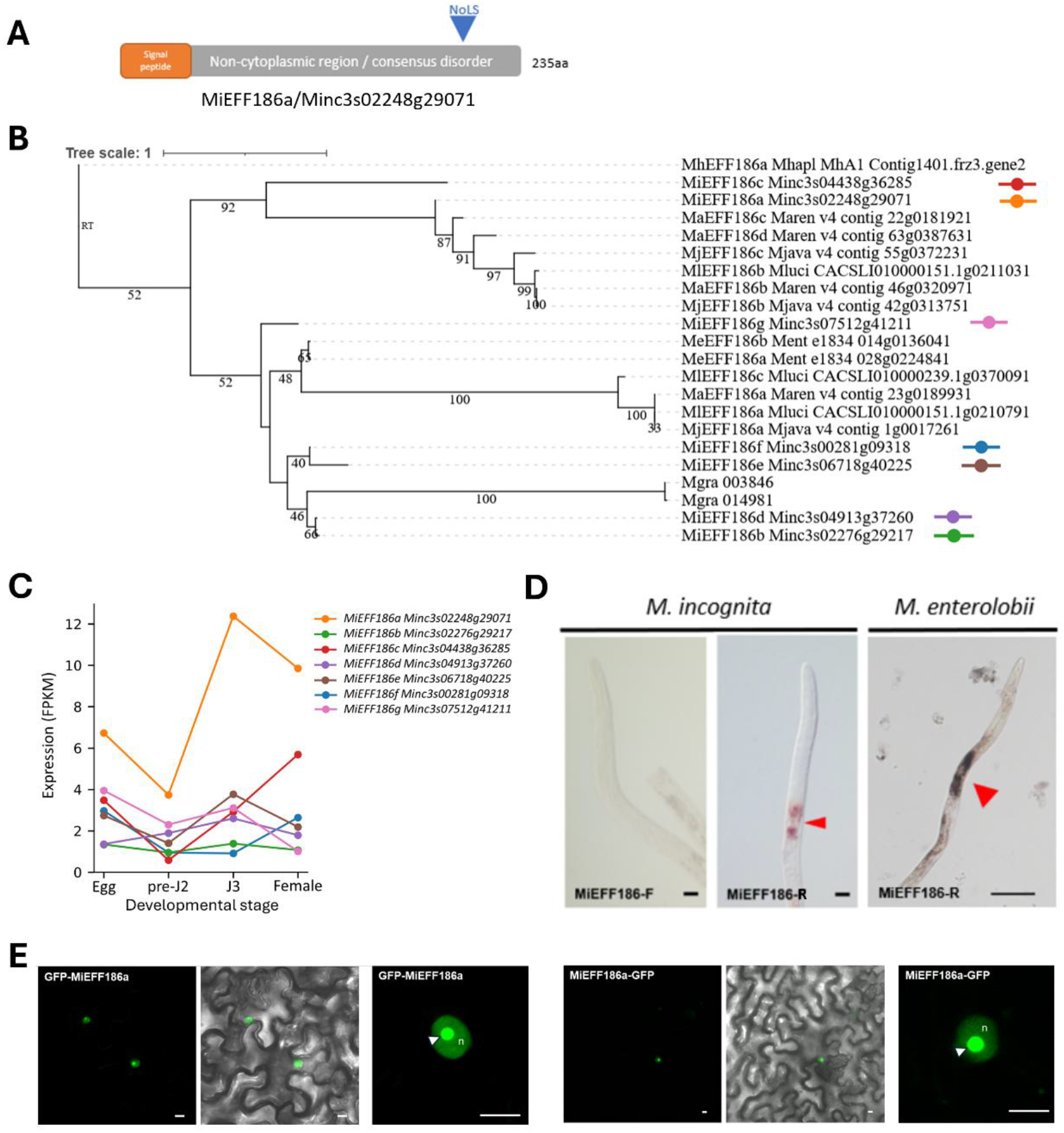
MiEFF186 is a conserved RKN secreted effector that targets the plant nucleus and nucleolus. **A,** Schematic diagram of the MiEFF186 protein. The protein includes a predicted secretion signal peptide (red box), a non-cytoplasmic region harboring a consensus disorder X (grey box) and a nucleolar localization signal (NoLS). **B,** Phylogenetic tree of *Meloidogyne* spp. EFF186 amino acid sequences. The number of tree replicates in which the associated taxa were assembled in 100 bootstraps is indicated. **C,** Transcript abundance of *MiEFF186* genes in *M. incognita* eggs, pre-parasitic juveniles (pre-J2), parasitic juveniles (J3) and females was retrieved from RNAseq data (Blanc-Mathieu *et al*., 2017) and is displayed as Fragments Per Kilobase per Million mapped fragments (FPKM). **D,** *In situ* hybridization, showing *EFF186* transcripts in the subventral glands (SvGs; arrowheads) of pre-parasitic juveniles (J2) of *M. incognita* and *M. enterolobii*. Images of J2s treated with sense (F) and anti-sense (R; negative control) *EFF186* probes are shown. Bars, 40 μm. **E,** MiEFF186 localizes in the nucleus (n) of plants cells and accumulates in the nucleolus (arrowhead) of plant cells. MiEFF186a coding sequence without SP was fused to the GFP in a N-or C-terminal position and expressed in *N. benthamiana* leaves by agroinfiltration. Bars = 10 µm.

*In-situ* hybridization experiments were performed on *M. incognita* and *M. enterolobii* pre-parasitic second stage juveniles (pre-J2) using a *MiEFF186a*-specific antisense probe. *MiEFF186* transcripts were shown to be synthesized specifically in the subventral esophageal glands of the nematodes (Fig 2D). No signal was detected in *M. incognita* pre-J2 when using a sense probe (Fig. 2D).

To determine the putative localization of MiEFF186 *in planta*, we infiltrated 3-week-old *N. benthamiana* leaves with *A. tumefaciens* strains carrying p35S::GFP-MiEFF186a or p35S::MiEFF186a-GFP constructs. Using both fusions, MiEFF186a localized in the nucleus and accumulated in the nucleolus, consistent with the presence of a NoLS (Fig. 2E). Taken together, these results suggest that MiEFF186a is secreted *in planta* during plant-RKN interaction and targets the plant cell nucleolus.

### 2.3 | MiEFF186a interacts with a host splicing factor

In order to identify tomato plant targets of MiEFF186a, a Y2H screen was performed by using MiEFF186a without its SP as a bait, and an infected tomato root cDNA library as the prey (Hybrigenics Services; Zhao *et al*., 2020; Mejias *et al*., 2021; Soulé *et al*., 2024). The screen of 110 million interactions identified six different putative targets (Fig. 3A). In addition to the peptides usually captured under our experimental conditions (Solyc09g076020.2 and Solyc07g053740.1) and those captured only once or twice, we identified one putative tomato target, encoded by *Solyc08g081580* which was captured five times. *Solyc08g081580* is orthologous to *Arabidopsis thaliana CRY2 INTERACTING SPLICING FACTOR2* (*CIS2*: AT1G63980; Zhao *et al*., 2022), also known as *G-PATCH DOMAIN PROTEIN1* (*GDP1*; Kojima *et al*., 2018). We investigated further the interaction between MiEFF186a and GDP1/CIS2 using several strategies. Firstly, the interaction between MiEFF186 and full length GDP1/CIS2 sequences has been confirmed by pairwise Y2H. Yeasts containing the bait plasmid MiEFF186 and the prey plasmids expressing *SlGDP1/CIS2* from *Solanum lycopersicum*, *AtGDP1* from *A. thaliana* or *NbGDP1b* from *N. benthamiana* were mated. Diploid yeasts have been plated on non-selective media depleted in tryptophan (W) and leucine (L), and selective media depleted in W and L and histidine (H), including 1 mM 3-amino-1,2,4-triazole (3-AT), a competitive inhibitor of the *HIS3* reporter gene (Fig. 3B). MiEFF186a interacts with SlGDP, AtGDP1 and NbGDP1 in yeast. No interaction was detected between MiEFF186a and the SV40 large T-antigen or between MiEFF16, another *M. incognita* nucleolar effector (Mejias *et al*., 2021), and GDP1s (Fig. 3B). Secondly, co-immunoprecipitation assays were conducted to validate the interaction between MiEFF186a and the tomato GDP1/CIS2. The full length was fused to the GFP to create a GFP:GDP1 construct. It was co-expressed together with MiEFF186a without its SP and fused to the RFP (RFP:MiEFF186), or with the RFP alone (DsRED), in *N. benthamian*a leaves (Fig. 3C). RFP:MiEFF186 could be co-immunoprecipitated with GFP:GDP1, but not the DsRED alone (Fig. 3C) Thirdly, to test whether MiEFF186a and the tomato GDP1/CIS2 might localize to the same cellular compartment, constructs using the fluorescent N-terminal tags RFP and GFP fused to the MiEFF186a without SP and SlGDP1/CIS2, respectively, were co-expressed into 3-week-old *N. benthamiana* epidermal leaf cells. Co-expression experiments confirmed the localization of RFP:MiEFF186 with GFP:GDP1 in the nucleolus (Fig. 3D). Unlike RFP:MiEFF186, GFP:GDP1 was exclusively detected in the nucleolus and was not observed in the nucleoplasm, consistent with previous observations (Kojiama *et al*., 2018). Interestingly, RFP:MiEFF186 and GFP:GFP1 appeared to co-localize at a ring-like structure at the inner periphery of the nucleolus (Fig. 3D). Taken together, our results demonstrate that MiEFF186 interacts with GDP1/CIS2 at the edge of the nucleolus.

**Figure 3.**
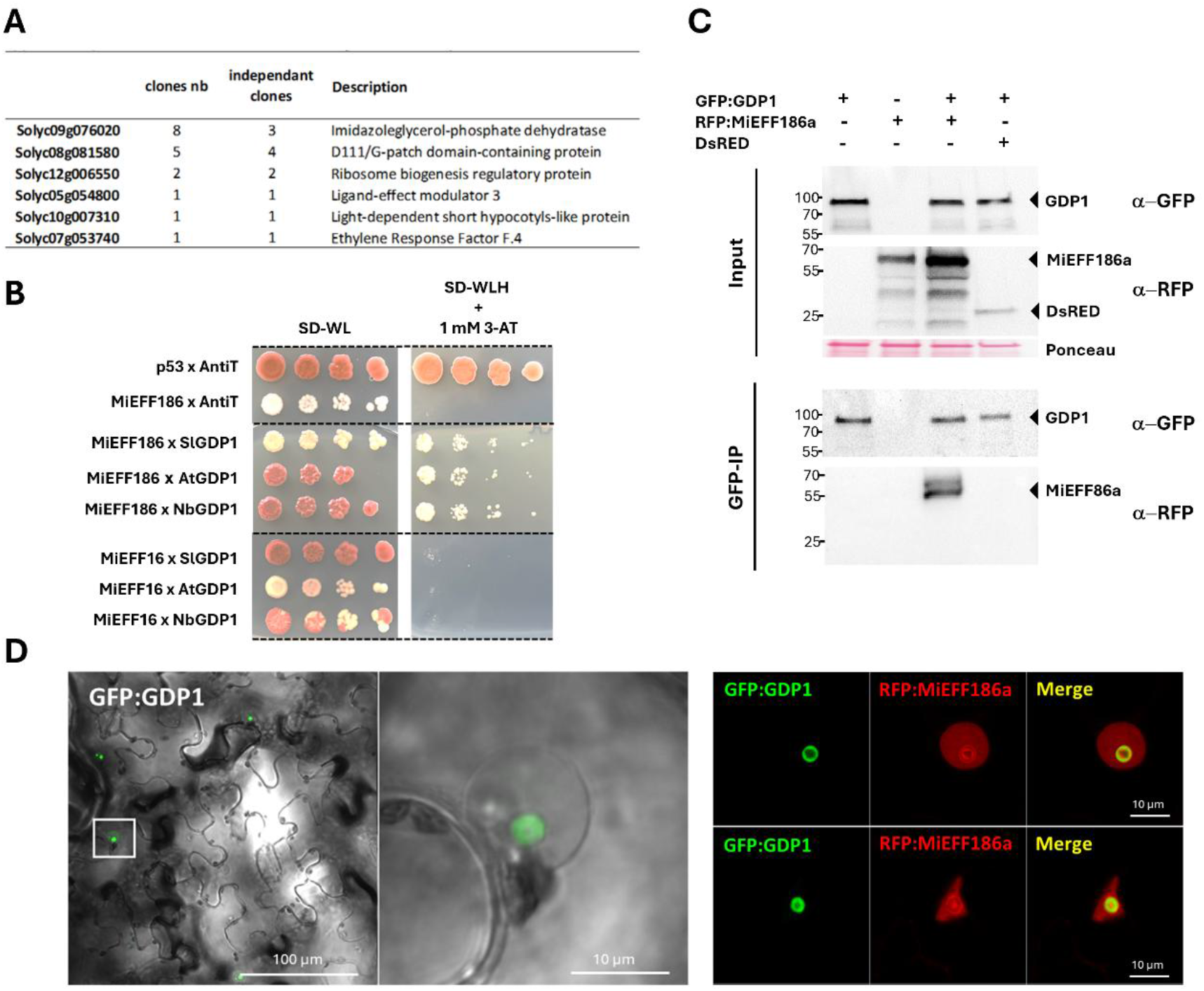
MiEFF186 interacts with GDP1 in the plant cell nucleolus. **A,** List of putative MiEFF186a-interacting tomato proteins identified in our Y2H screen. **B,** Diploid yeasts containing the bait and prey plasmids carrying controls, effectors or GDP1 proteins (*Solanum lycopersicum* SlSmD1 and *Arabidopsis thaliana*, AtGDP1 and *N. benthamiana*, NbGDP1) were spotted on plates. SD-WL corresponds to the nonselective medium without tryptophan (W) and leucine (L). Only yeasts carrying a protein–protein interaction can survive on the SD-WLH (H, histidine) + 1M 3-AT selective medium. Murine p53 and SV40 T-antigen T (anti T) were used as a positive control, and MiEFF16 was used as a negative control. **C,** Co-immunoprecipitation (Co-IP) experiments confirmed that MiEFF186a interacts with the tomato SlGDP1. GFP:GDP1 was transiently co-expressed with RFP:MiEFF186a or DsRED alone in *N. benthamiana* leaves. The Co-IP experiment was performed using anti-GFP affinity beads, and the isolated proteins were analyzed by western blotting with anti-GFP antibodies to detect SlGDP1, and with anti-RFP antibodies to detect MiEFF186a or DsRED. **D,** Single-plane confocal images of *N. benthamiana* leaf epidermal cells infiltrated with *Agrobacterium tumefaciens* and producing SlGDP1, fused to the C-terminal end of the green fluorescent protein (GFP) reporter (GFP:GDP1; green signal) (left panel), and producing both GFP:SlGDP1 (GFP:GDP1) and MiEFF186a fused to the C-terminal end of the red fluorescent protein (RFP) reporter (RFP:MiEFF12a; red signal) (right panel) ;Overlays of fluorescence images are shown (merge).

### 2.4 | CIS2/GDP1 is involved in plant susceptibility to RKNs

We then investigated the potential function of GDP1/CIS2 in RKN parasitism, by using two previously described T-DNA homozygous knockout mutant lines for *GDP1/CIS2*, i.e. *gdp1-1* and *gdp1-2*, and a complemented line (Kojima *et al*., 2018). These mutant lines were inoculated with 200 pre-J2 of *M. incognita* and gall number was evaluated in mutant roots as compared to the complemented line pGDP1::GDP1-GUS and wild-type line (Col-0) (Fig. 4A). Six weeks after inoculation, the number of galls was significantly reduced, as compared to the wild-type or the complemented line (Fig. 4A). We then designed a chimeric TRV2 construct to silence *NbGDP1/CIS2* genes. This construct specifically targeted a 233 and 234 nucleotides region of NbGDP1a and NbGDP1b, respectively (Supplementary Fig. S6 and S7). RT-qPCR validated silencing of GDP1/CIS2 genes by 71% in *N. benthamiana* 10 days post-inoculation with the TRV (Fig. S7). 10 days post-inoculation with the TRV, plants were inoculated with 200 pre-J2. Five weeks later, the number of egg masses found on the roots of the silenced plants was significantly reduced when compared to the TRV2-GFP control (Fig. 4B). These results demonstrate an important role for CIS2/GDP1 in plant susceptibility to *M. incognita*.

**Figure 4.**
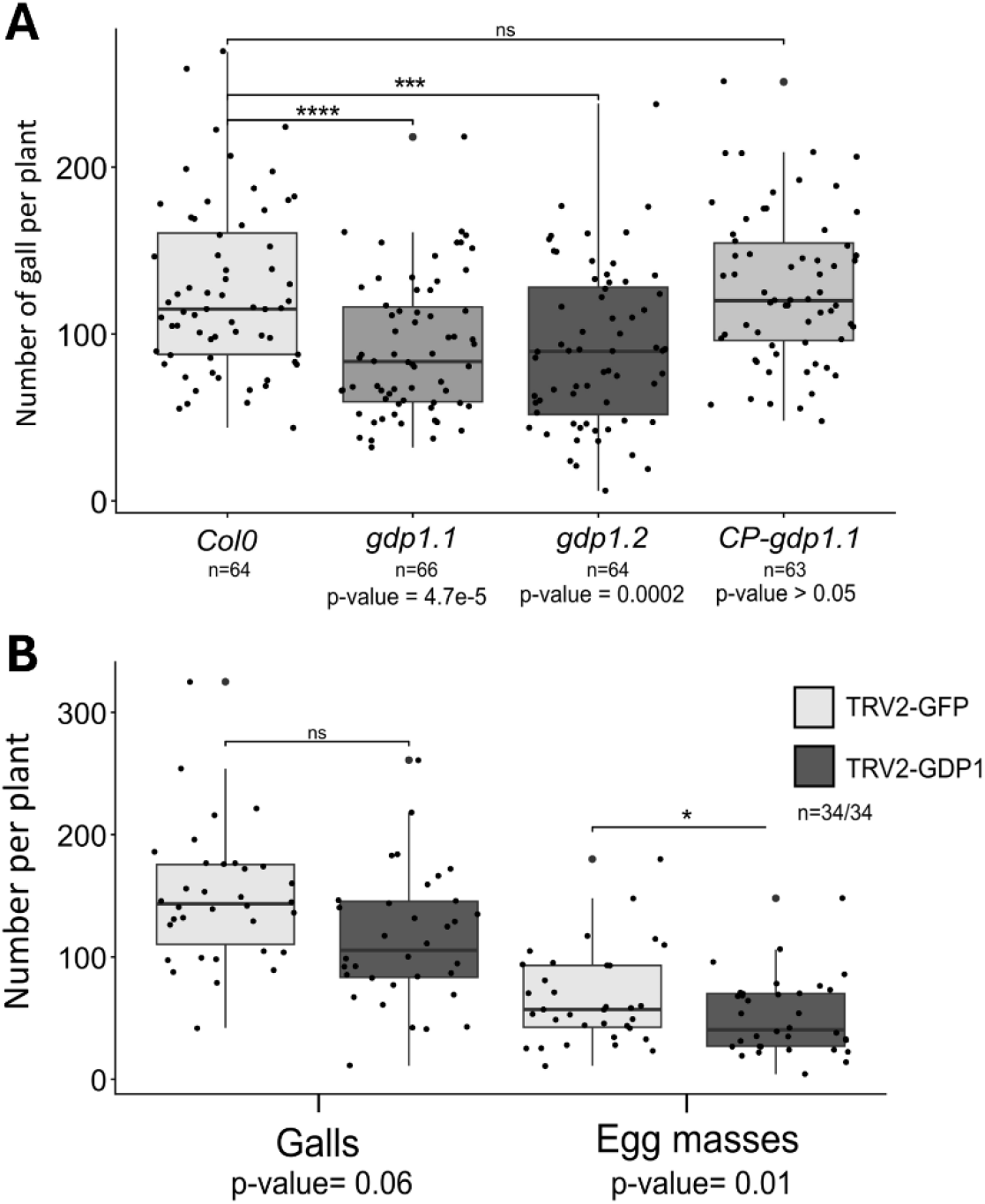
GDP1/CIS2 contributes to *Arabidopsis* and *N. benthamiana* susceptibility to *M. incognita*. **A,** Box-and-whisker plots of galls per plant in Col-0 control line, *gdp1.1* and *gdp1.2* mutants and a pGDP1::GDP1-GFP/gdp1-1 complemented line (CP-gdp1.1) six weeks post infection with 200 *M.incognita* second-stage juveniles (J2s). Box indicates interquartile range (25th to the 75th percentile). The central line within the box represents the median. Whiskers indicate the minimum and maximum values for the normal values present in the dataset. Statistical significance was assessed using Wilcoxon rank sum test with continuity correction (***p<0.001 ; ***p<0.0001). **B,** Infection test on *N. benthamiana* TRV-GFP or TRV2-GDP1s plants. Galls and egg masses were counted five weeks after inoculation with 200 *M. incognita* second-stage juveniles (J2s) per plant. Box indicates interquartile range (25th to the 75th percentile). The central line within the box represents the median. Statistical significance was assessed using Wilcoxon rank sum test with continuity correction (*p≤0.01).

### 2.5 | *MiEFF186* and *AtGDP1/CIS2* affect gene expression and AS in *Arabidopsis* roots

Since GDP1/CIS2 has been shown to encode a splicing factor in *Arabidopsis* (Zhao *et al*., 2022), we first examined whether this function was conserved in tomato. We performed RT-qPCR to verify that expression of our *GFP:GDP1* construct could affect AS by quantifying *RLPK.1* and *RLPK.2* transcripts in the *35S::RLPK-LUC* transgenic *N. benthamiana*, and we demonstrated a significant effect of *SlCIS2/GDP1* expression in modulating AS in the reporter system (Supplementary Fig. S2).

To investigate the effect of MiEFF186a secretion and the effect of its interaction with the host GDP1/CIS2 on plant transcriptome and on modulating AS, we analysed at a genome-scale level the *gdp1-2* mutant line and in a transgenic *Arabidopsis* line overexpressing *MiEFF186a* (MiEFF186-OE), compared to the wild-type Col-0. We performed RNA-sequencing (RNA-seq) on total RNA isolated from the roots of plantlets cultured in liquid medium, as described previously (Mejias *et al*., 2021; Soulé *et al*., 2024). The RNA seq data analysis identified 86 and 372 differentially expressed genes (DEG) in the MiEFF186-OE and the *gdp1.2* mutant, when compared to Col-0, respectively (FDR < 0.05, |log2FC| > 1) (Fig. 5A and 5B). Noteworthy, one third of the DEG in the MiEFF186a-overexpressing line were all differentially expressed in the gdp1.2 mutant (Fig. 5B). Gene ontology (GO) enrichment analysis was performed on DEG, induced or repressed genes, of each line. Significant terms were found only among genes induced in gdp1-2. A significant enrichment in terms in biological processes related to response to stress such as “response to stress”, “response to external stimulus” or “response to biotic stimulus” was observed among genes induced in the *gdp1.2* mutant (Fig. 5C). We then performed a SUPPA2 analysis to characterize splicing events and quantify each transcripts isoforms in the *gdp1.2* mutant and the MiEFF186a-overexpressing line. We identified 466 and 257 differential splicing events in the *gdp1.2* mutant and the *MiEFF186a*-overexpressing line, and when compared to Col-0, respectively (Supplementary Fig. S8). Intron retention, exon skipping, and alternative 5’/3’ splice site selection events affected 316 and 171 genes, in the *gdp1.2* mutant and the *MiEFF186a*-expressing line, respectively (Fig.5D). Most of the splicing events in the *gdp1.2* and the *MiEFF186a*-overexpressing lines are intron retention and selection of alternative 3’ splice site (Supplementary Fig. S8). A high degree of overlap was observed between the two lines, with almost a half (72/171 genes) of the differentially spliced genes in the *MiEFF186a*-overexpressing line being also differentially spliced in the *gdp1.2* mutant (Fig. 5D), Thus, MiEFF186 and its target GDP1/CIS2 might participate to AS triggered in *Arabidopsis* following infection with *M. incognita* to modulate host transcriptome and immunity.

**Figure 5.**
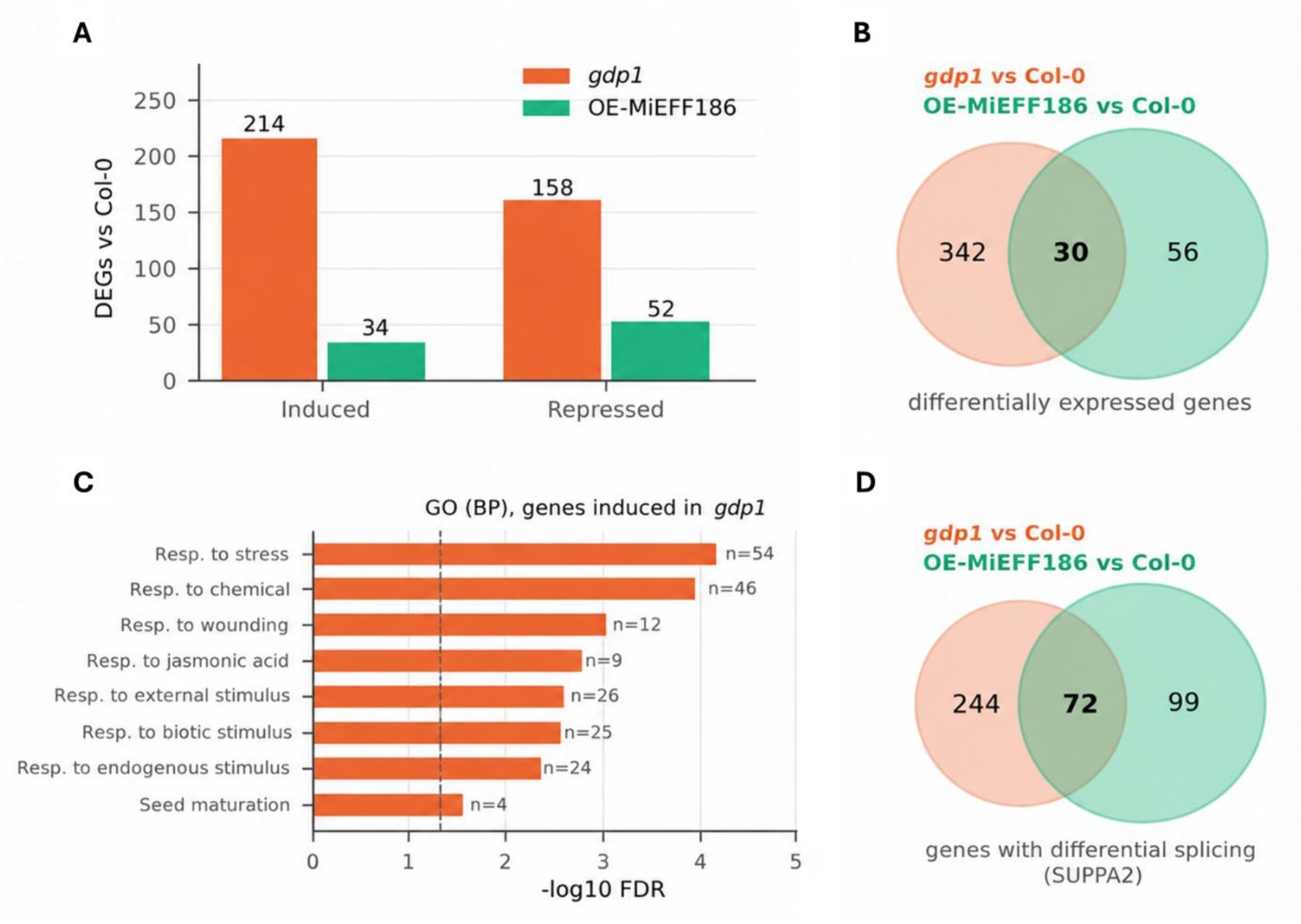
MiEFF186a and GDP1/CIS2 regulate transcript accumulation and alternative splicing in Arabidopsis root. **A,** Quantification of differentially expressed genes (DEG), up- or down-regulated, in the roots of the MiEFF186-expressing Arabidopsis line (EFF18) and the gdp1.2 mutant, relative to Col-0 roots. **B,** Venn diagram showing the overlaps between differentially expressed genes in the MiEFF186-expressing line, and the *gdp1.2* mutant. **C,** GO enrichment analysis on the 214 genes upregulated in the gdp1.2 mutant indicating an enrichment in genes related to the stress responses. **D**, Venn diagram showing the overlaps between differentially spliced genes in the MiEFF186a-expressing line, and the *gdp1.2* mutant.

## 3 | DISCUSSION

Pre-mRNA splicing and AS are important post-transcriptional processes that regulates plant growth and development as well as responses to biotic and abiotic stresses. Plant immune responses are modulated through AS regulation, and pathogens are known to manipulate AS for successful parasitism (Rigo *et al*., 2019; Hewezi, 2024). Consistently, several effectors from pathogens have been shown to target key components of the AS machinery, including RNA-binding splicing factors (Hewezi, 2024; Godinho *et al*., 2025; Gao *et al*., 2026). For example, the bacterial effector HopU1 from *Pseudomonas syringae* interacts with several RNA-binding proteins, including GRP7, in Arabidopsis. Through its ribosyltransferase activity, HopU1 modifies the RNA-binding domain of GRP7 preventing its binding to transcripts involved in the host immunity (Nicaise *et al*., 2013). The *Phytophthora sojae* effector PsAvr3c interacts with and stabilizes two soybean serine/lysine/arginine-rich proteins (GmSKRPs), which interact with various proteins of the spliceosome to regulate splicing and host immunity (Huang *et al*., 2017). Another example is the PsFYVE1 effector which modulates AS of immunity-related genes by interacting with the RNA-binding protein RZ-1A in *N. benthamiana* (Lu *et al*., 2023). Plant parasitic nematodes secrete such SREs, such as the *Heterodera schachtii* Hs30D08 effector that binds to the *Arabidopsis* auxiliary spliceosomal protein SMU2 and the RKN EFF18 that target plant SmD1 (Verma *et al*., 2018; Mejias *et al*., 2021 and 2022). A luciferase-based splicing reporter using transgenic *N. benthamiana* expressing a RLPK-luciferase reporter was initially developed to identify *P. infestans* SREs (Huang *et al*., 2020). Subsequently, this reporter system has been used successfully to identify SREs from the bacteria *Ralstonia solanacearum* and the fungus *Exserohilum turcicum* (Li & Kou, 2025; Yu *et al*., 2025). Here, using the same approach, we identified MiEFF186a as a new RKN SRE. *EFF186* genes are exclusive to RKN but conserved among RKN species. EFF186 putative effectors have no predicted functional domain but own a nucleolar localization signal. Consistently, MiEFF186a effector localized exclusively in the nucleus and accumulated in the nucleolus when transiently expressed in *N. benthamiana* leaf epidermal cells. Only a very few plant pathogen effectors have been shown to target this nuclear body which is the site of key functions such as rRNA synthesis, ribosome biogenesis and assembly of ribonucleoproteins such as spliceosomal snRNPs (Lee *et al*., 2024; Ranty-Roby *et al*., 2024).

Using a yeast two hybrid approach, we demonstrate that MiEFF186a interacts with the nucleolar protein GDP1/CIS2 of tomato, *Arabidopsis* and *N. benthamiana*. GDP1/CIS2 contains a G-patch domain generally found in RNA-binding proteins involved in RNA processing, such as pre-mRNA splicing and rRNA and ribosome biogenesis (Karika *et al*., 2026). In *Arabidopsis thaliana*, both *CRYPTOCHROME-INTERACTING SPLICING FACTOR* (*CIS1*) and its paralogue gene *CIS2*, regulate AS of the *FLOWERING LOCUS M* gene (*FLM*) and thermosensory flowering (Zhao *et al*., 2022). Remarkably, in *Arabidopsis*, CIS1’s activity in modulating *FLM* AS requires its interaction with the large subunit,

U2AF65, of the U2 snRNP auxiliary factor (U2AF) (Zhao *et al*., 2022). Another RKN effector, MiMSP8, was shown to interact with the tomato SlU2AF35, the small subunit of the U2AF heterodimeric splicing factor, impeding its association with a subset of pre-mRNAs (Chen *et al*., 2025). Thus, CIS1/CIS2 and U2AF could be part of a modular hub on which different RKN effectors converge. Interestingly, this U2AF splicing factor is targeted upon diverse biotic interactions, such as those involving pathogenic virus or symbiotic arbuscular mycorrhizal fungi (Du *et al*., 2024; Betz *et al*., 2024). Consistently, MiEFF186a modified splicing of a reporter transcript. In addition, *gdp1* and *MiEFF186a*- overexpressing roots shared part of their splicing and expression changes, and genes induced in *gdp1* were enriched in defence-related functions. These changes may reflect a direct role of GDP1/CIS2 in splicing.

In *Arabidopsis*, GDP1 was also described for its role in ribosome biogenesis. GDP1 was shown to be required for pre-rRNA processing and maturation (Kojima *et al*., 2018). Interestingly, a second putative tomato target of MiEFF186a identified in this study (Solyc12g006550, capture only twice in our Y2H screen; Fig. 3A) is a homologue of the *Arabidopsis* RIBOSOME BIOGENESIS REGULATOR1 (ARRS1) a nucleolar ribosome biogenesis factor required for pre-rRNA processing and the biogenesis of the 60S ribosomal subunit (Choi *et al*., 2020; Maekawa *et al*., 2018). Altering ARRS1 levels in Arabidopsis affects leaf morphology and plant lifespan, indicating that this pathway influences plant growth (Maekawa et al., 2018). Similarly, GDP1/CIS2 promotes leaf cell proliferation through its role in ribosome biogenesis (Kojima et al., 2018). Moreover, MiEFF186a and GDP1/CIS2 co-localised at the nucleolar periphery, a region associated with early ribosome assembly. Ribosome biogenesis is targeted by diverse plant pathogens (Lee et al., 2023; Ranty-Roby et al., 2024).The *Aphanomyces euteiches* effector AeSSP1256 similarly relocates the RNA helicase MtRH10 to the perinucleolar space and alters its function in ribosome biogenesis (Camborde et al., 2022). In plant-parasitic nematodes, the cyst nematode effector 32E03 also manipulates this pathway, by targeting histone deacetylases, it alters rDNA chromatine structure and rDNA expression and regulate pre-rRNA abundance (Vijayapalani *et al*., 2018). Since giant cells are hypertrophied and metabolically hyperactive, MiEFF186a may target ribosome biogenesis to adapt protein synthesis to the demands of the feeding site.

The functions of GDP1/CIS2 require further investigation, particularly by identifying its targets in the context of the plant-RKN interaction. We demonstrated that GDP1/CIS2 plays a crucial role in facilitating successful root-knot nematode (RKN) infection in both Arabidopsis and members of the Solanaceae family. *GDP1/CIS2* is therefore an interesting susceptibility gene that could be targeted to engineer durable resistance to RKNs.

## 4 | EXPERIMENTAL PROCEDURES

### 4.1 | Nematode and plant materials

*M. incognita* strain ‘Morelos’ was multiplied on tomato (*Solanum lycopersicum* cv ‘Saint Pierre’) grown in a growth chamber (25°C, 16 h : 8 h, light : dark photoperiod). Freshly hatched second-stage juveniles (J2s) were collected as described previously (Caillaud & Favery, 2016). *Arabidopsis thaliana gdp1.1* and *gdp1.2* mutants and the complemented line have been described previously (Kojima et al., 2018). For RKN infection, plants were grown with Columbia 0 accession plants in sand:soil (50:50) in small squared pots in short day conditions (8h light/16h dark), 22°C for four weeks then were inoculated with RKN J2s.

For RNA-seq analysis, Arabidopsis were grown in liquid MS medium (½MS salts, 1% sucrose, pH 6.4) with gentle shaking (70 rpm), under a 12 h:12h, light:dark photoperiod, at 25°C, as described previously (Mejias *et al*., 2021). Roots were collected after 11 d and immediately frozen in liquid nitrogen until RNA extraction.

For splicing reporter assays, transgenic *N. benthamiana* carrying a p35S::RLPK-LUC construct (Huang *et al*., 2020) were grown in soil at 24°C under long-day conditions (16h light/8h dark). For VIGS, *N. benthamiana* seeds were sown in soil and grown at 24°C under long-day conditions (16h light/8h dark). Seedlings were transplanted into individual pots containing the sand:soil mixture (50:50) after cotyledon emergence. Plants were maintained at 24°C until further treatments. For agro-infiltration of co-localization constructs, *N. benthamiana* wild type plants were grown in soil for 3 weeks before agro-infiltration.

### 4.2 | Screening for effectors involved in alternative splicing, and validation by RT-qPCR

A splicing reporter system already reported to screen for the effectors involved in splicing regulation (Huang et al., 2020) has been used. Leaves of transgenic *N. benthamiana* lines carrying a RLPK-LUC construct, were agroinfiltrated with *A. tumefaciens* containing GFP-effector fusions. Then plants were treated with 1 mM luciferin. Then, leaf discs were collected 48 hours after infiltration and LUC activity was measured using a luminometer. All data were analyzed for significant differences (Kruskal-Wallis test and post hoc Dunn-Bonferroni test; adjusted p-value<0,01)

To quantitatively test the alternative splicing activity of nematode effectors and tomato genes, six-week-old 35S-RLPK-LUC transgenic *N. benthamiana* leaves were infiltrated with *A. tumefaciens* strains containing either a GFP-effector/gene plasmid or a GFP control plasmid. After 48 hours, the infiltrated leaves were collected for observation of gene expression under a confocal microscope (LSM880, Zeiss, Germany). Leaves exhibiting strong GFP signals were selected for RNA extraction. Total RNA was isolated using the RNeasy Plant Mini Kit (Qiagen, Germany) and quantified with a NanoDrop spectrophotometer (Thermo Fisher, USA). One microgram of total RNA was used for cDNA synthesis with the PrimeScript™ RT Reagent Kit (TAKARA, Japan). RT-qPCR was carried out using the Brilliant III SYBR Master Mix (Agilent, USA) on a CFX Connect Real-Time Quantitative PCR system (Bio-Rad, USA). The *NbGAPDH* and *NbEF1α* genes (Supplementary Table S1) were used as reference genes to normalize gene expression using the 2−ΔΔCt method. Alternative splicing events were validated by measuring the transcript levels of *RLPK.1* and *RLPK.2* using specific primers (Huang *et al*., 2020; Supplementary Table S1). Then, the splicing ratio was calculated. All RT-qPCR experiments were performed in three technical replicates and three biological replicates.

### 4.3 | EFF186 sequence analysis, alignment, and phylogenetic tree

The sequences of MiEFF186a putative paralogs and orthologs were obtained from Meloidogyne genomic resources (http://www6.inra.fr/meloidogyne_incognita) and Wormbase parasite (https://parasite.wormbase.org/index.html). Protein sequences were aligned with the MAFFT tool on the EBI server (https://www.ebi.ac.uk/Tools/msa/mafft/). The alignment was then used as input to build the maximum likelihood phylogenetic tree using IQTree (https://www.hiv.lanl.gov/content/sequence/IQTREE/iqtree.html). Support for the nodes was calculated with 100 bootstrap replicates. *Meloidogyne hapla* was used as the outgroup in the phylogenetic tree. The tree was visualized in iTOL https://itol.embl.de/. The pairwise sequence identity matrix of RKN EFF186 protein sequences was generated with Sequence Demarcation Tool version 1.2 software (Muhire et al., 2014) (http://web.cbio.uct.ac.za/∼brejnev/).

### 4.4 | *In situ* hybridization

Approximately 10,000 J2s of M. incognita and *M. enterolobii* were collected and *in situ* hybridization was performed as previously described Jaouannet *et al*. (2018). For probe production, *MiEFF186a* sequence was amplified using M. incognita J2 cDNAs. Sense probes were used as negative controls. Pictures were obtained using Axioplan2 (Zeiss) microscope.

### 4.5 | Molecular cloning

The coding sequence (CDS) of *M. incognita* MiEFF186a without the signal peptide (SP), along with *Arabidopsis*, *S. lycopersicum* and *N. benthamiana CIS2/GDP1* coding sequences were amplified using PCR with specific primers (Supplementary Table S1). These PCR products were inserted into the pDON207 donor vector and recombined into Pp6 (Y2H prey vector), pK7WGR2 (RFP:MiEFF186a), pK7GWF2 (GFP:SlGDP1/CIS2; GFP:MiEFF186a), and pK7FWG2 (MiEFF186a:GFP) using Gateway technology (Invitrogen), as described by Karimi et al., 2002. Nb-GDP1 VIGS best region has been identified using the SGN VIGS Tool for both genes (Fernandez-Pozo et al., 2015) and chimeric TRV2 construct containing both regions was synthesized into a pUC vector (Azenta Life Sciences, Leipzig, Germany ; Supplementary Fig. S7) and introduced in the TRV2 plasmid. All constructs were sequenced (GATC Biotech) and subsequently transformed into yeast Y187 or A. tumefaciens GV3101.

### 4.6 | Fluorescence microscopy

For transient expression in *N. benthamiana* leaves, 3-to 4-week-old *N. benthamiana* plants were agro-infiltrated using recombinant *A. tumefaciens* strains harboring GFP or RFP vectors as described previously (Soulé *et al*., 2024). Imaging was performed on leaf discs 48-hours post-infiltration with a Zeiss LSM 880 inverted confocal microscope; Samples were excited at 488 nm for GFP and 543 nm for RFP. GFP and RFP emissions were detected selectively with 505–530 and 560–615 nm band-pass emission filters respectively.

### 4.7 | Yeast two hybrid

For the yeast two-hybrid (Y2H) screening, the CDS of MiEFF186, excluding the SP, was employed as the bait. The MiEFF186a sequence was amplified (Supplementary Table S1) and cloned into the pB27 vector (Hybrigenics Services, Paris, France). The Y2H screening was carried out using a cDNA library derived from infested tomato roots (Hybrigenics Services, Paris, France), as described previously (Zhao *et al*., 2020; Mejias *et al*.,2021; Soulé *et al*., 2024).

For pairwise Y2H assays, full-length controls, baits, and potential targets (MiEFF18 and MiEFF16 without the SP, SlGDP1, AtGDP1, NbGDP1b and Antigen T) were cloned into either the pB27 or pP6 vector (Hybrigenics Services, Paris, France). The constructs were confirmed via sequencing and used to transform the L40ΔGal4 (MATa) or Y187 (MATα) yeast strains. Following mating between the Y187 and L40ΔGal4 strains, diploid cells were selected on a medium lacking tryptophan and leucine. Interaction testing was conducted on a medium deficient in tryptophan, leucine, and histidine, with the addition of 1 mM 3-amino-1,2,4-triazole (3-AT).

### 4.8 | Infection assays

Four-week-old Arabidopsis seedlings or *N. benthamiana* plants infested with the TRV were inoculated with 200 M. incognita J2s per plant. Roots were harvested five weeks post-infection and stained with 0.5% eosin. Gall and/or the number of females producing egg masses sand the root weight were measured. Three independent biological replicates were conducted for each experimental condition. Graphs and plots were created with R and Microsoft® Office Excel® 2019. Statistical analyses were performed with R software (R Development Core Team, version 4.1.0).

### 4.9 | Virus induced gene silencing (VIGS)

The following *Agrobacterium tumefaciens* strains were used for agroinfiltration and VIGS: TRV1, TRV2-GFP and TRV2-empty (controls), and TRV2-NbGDP1/CIS2 as described in Mejias *et al*. (2022). Approximately 20-25 plants were infiltrated per condition. 10 days post-agroinfiltration, root samples were collected from three plants for RNA extraction and RT-qPCR analysis as described above and plants were inoculated with 200 *M. incognita* pre-J2. Five weeks post-nematode inoculation, roots were washed and the number of galls and egg masses was counted as described above.

### 4.10 | Co-immunoprecipitation assays (Co-IP)

Immunoblots were performed using 30 µg of total protein separated on SDS-PAGE gels (4–15% Mini-PROTEAN® TGX™ Precast Protein Gels, Bio-Rad). For GFP detection, a mouse anti-GFP antibody (1:5000 dilution; Roche) was used. A mouse anti-RFP antibody (Chromotek; 1:5000 dilution) was used for RFP detection. An HRP-conjugated anti-mouse secondary antibody (Bio-Rad) was used with an ECL Clarity Kit (Bio-Rad) for visualization. Co-immunoprecipitation was performed as previously described (Camborde *et al*., 2022). *N. benthamiana* leaf samples were harvested 24 h after agroinfiltration, and total proteins (50 µg) were incubated for 3 h at 4°C with 30 µl of GFP-Trap Agarose beads (Chromotek, Planegg, Germany). After washing, the GFP-bound beads were boiled in SDS loading buffer (Merck) and immunoblotting was performed.

### 4.11 | Transcriptome analysis

For RNA-seq experiments, seeds were surface-sterilized and sown in liquid MS medium (½MS salts, 1% sucrose, pH 6.4) with gentle shaking (70 rpm), under a 12 h : 12 h, light : dark photoperiod, at 25°C. Roots were collected after 11 days and immediately frozen in liquid nitrogen. RNA was extracted using RNeasy plant mini kit (Qiagen). The RNA was treated with DNAse treatment using TURBO DNAse-free (Thermofisher), and its quality and integrity were assessed with Nanodrop. Library construction and paired-end sequencing were performed at the BGI (China). Differential gene expression was analysed using DESeq2 (FDR < 0.05, |log2FC| > 1) and differential alternative splicing using SUPPA2 (|dPSI| > 0.1), as described previously (Mejias *et al*., 2021). Gene Ontology enrichment analysis was performed as follows. Biological Process annotations were obtained from TAIR (ATH_GO_GOSLIM.txt TAIR release of 1 September 2026). Enrichment was tested for induced and repressed genes of each line, at the level of individual GO terms (5–500 annotated genes) and of TAIR GO-slim categories, using a one-sided hypergeometric test. All genes tested by DESeq2 were used as the background. P-values were corrected with the Benjamini–Hochberg procedure, and terms with FDR < 0.05 and at least two genes were considered enriched. Gene lists, overlaps, enrichment tests and plots were produced in Python (pandas, SciPy, Matplotlib).

## ACKNOWLEDGMENTS

We thank Prof. Gorou Horiguchi (Rikkyo University, Japan) for the kind gift of the Arabidopsis *gdp1.1 and gdp1.2* mutants and the pGDP1::GDP1-GFP/gdp1-1 complemented line. We thank Prof. Suomeng Dong (Nanjing Agricultural University, China) for the generous gift of the p35S::RLPK-LUC transgenic *N. benthamiana* and SRE3 and SRE4 constructs. We thank Dr Johnathan Dalzell and Steven Dyer (Queen’s University Belfast, UK) for VIGS protocol and vectors, Pr S.P. Dinesh-Kumar (University of California, Davis, USA) for VIGS vectors and Hybrigenics Services (Paris, France) for providing the pB27 and pP6 vectors and the L40ΔGal4 and Y187 yeast strains. Microscopy work was performed at the SPIBOC imaging facility of Institut Sophia Agrobiotech and we thank Dr Olivier Pierre for technical support. SRR holds a doctoral fellowship from Syngenta. This work is supported by INRAE and the French Government (National Research Agency, ANR) through the “Investments for the Future” LabEx SIGNALIFE (#ANR-11-LABX-0028-01) and the MASH project (#ANR-21-CE20-0002), the INRAE Syngenta Nem-targetome project, and the French-Chinese bilateral collaboration programs PHC XU GUANGQI 2020 #45478PF and PHC CAI YUANPEI 2024 #51126ZC.

## AUTHOR CONTRIBUTIONS

SRR, LC, JZ, MDC, JB, BF, MQ conceived and designed the experiments. SRR, KM, SS, LC, KH, JP, PL, CC, PF, MBL, CHC, JZ, JB, MQ performed the experiments. SRR, LC, SJ, JZ, MDC, JB, BF, MQ analyzed the data. JZ, BF, MQ supervised parts of the work. SJ, MDC, JB, BF, MQ contributed funding acquisition. SRR, BF, MQ wrote the article with input from all authors.

## CONFLICT OF INTEREST STATEMENT

The authors have no conflict of interest to declare.

## DATA AVAILABILITY STATEMENT

The data supporting the findings of this study are available from the corresponding author upon reasonable request.

---

**Supplementary Fig. S1.**
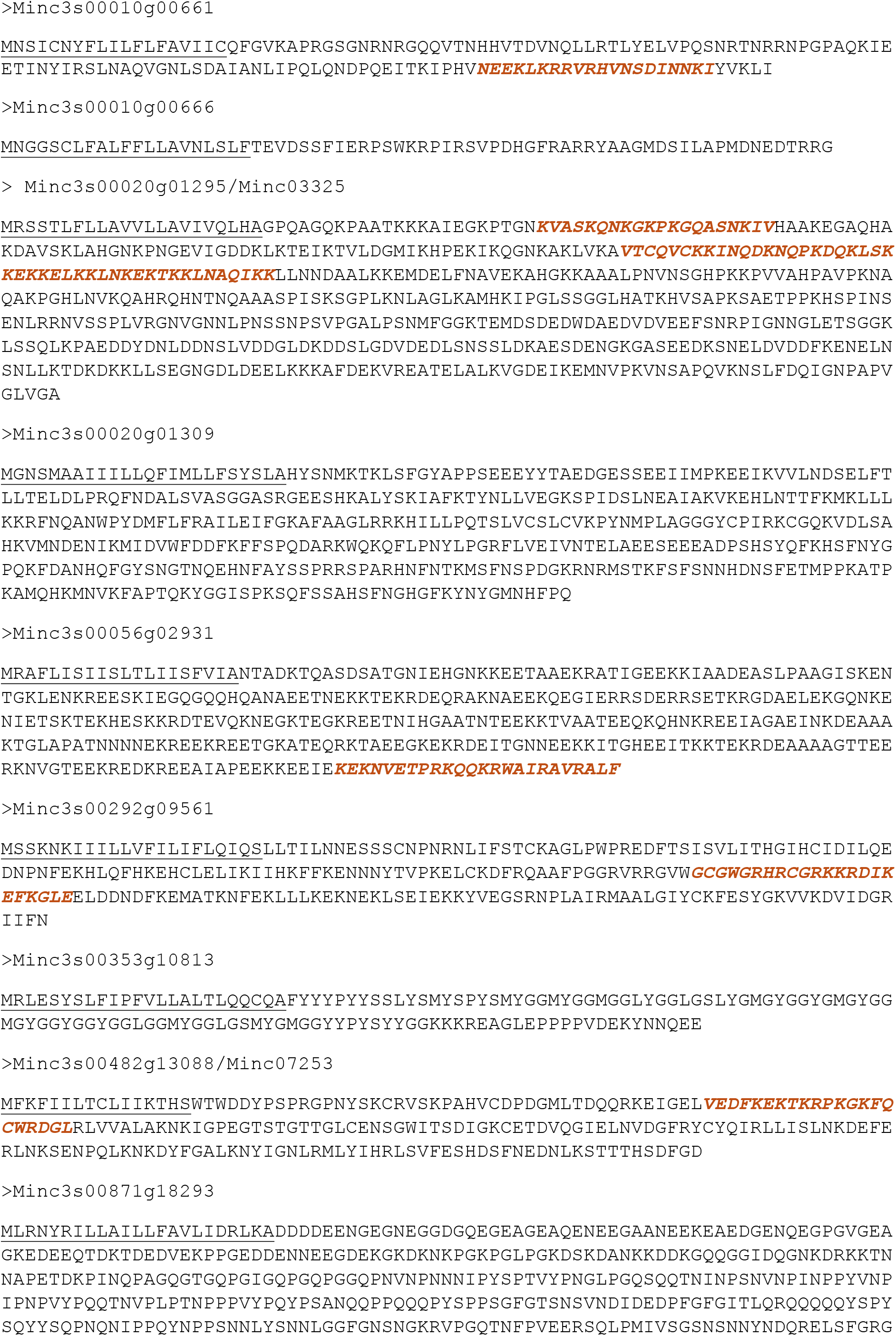
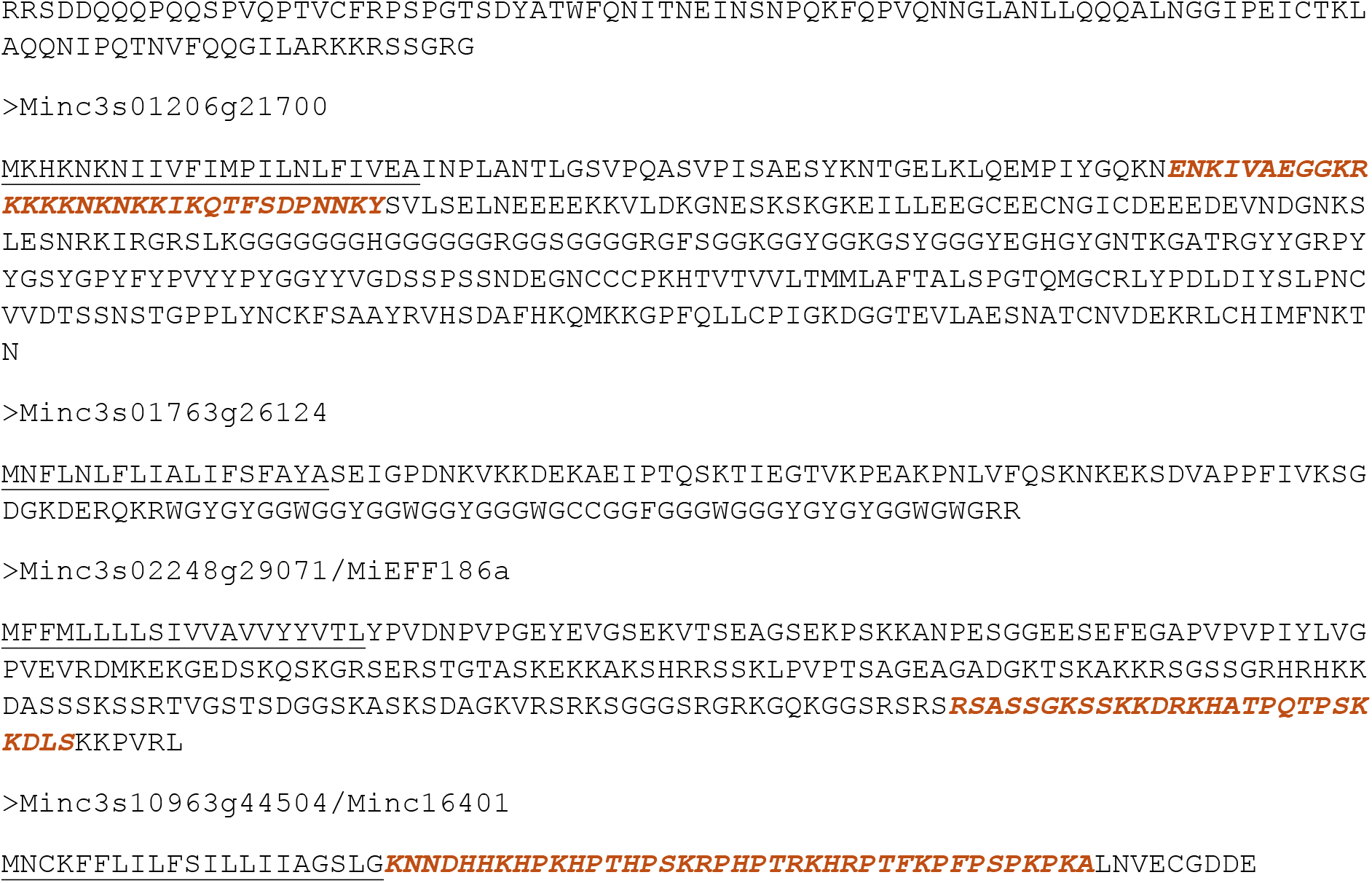
Amino acid sequences of effector proteins used in the splicing reporter assay. aa, amino acids. The signal peptide for secretion is underlined. Nucleolar Localisation Signal (NoLS) sequences are indicated in italics and orange.

**Supplementary Fig. S2.**
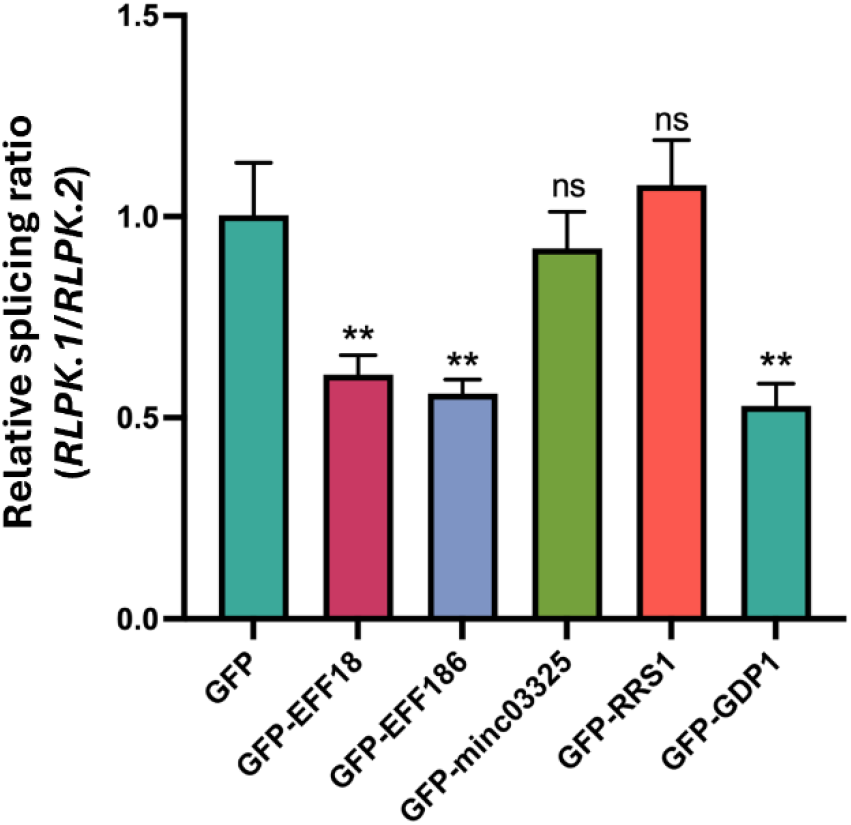
Validation of alternative splicing activity of RKN effectors and tomato genes by RT-qPCR. RT-qPCR experiments were performed to detect the relative expression levels of *RLPK.1* and *RLPK.2* of 35S-RLPK-LUC transgenic *N. benthamiana*. Leaves were infiltrated with *A. tumefaciens* strains containing a GFP-effector or -plant fusion or GFP control plasmid. *P*-values were from one-way ANOVA (ns, no significant difference; **, *P* < 0.01). This experiment was repeated three times with similar results.

**Supplementary Fig. S3.**
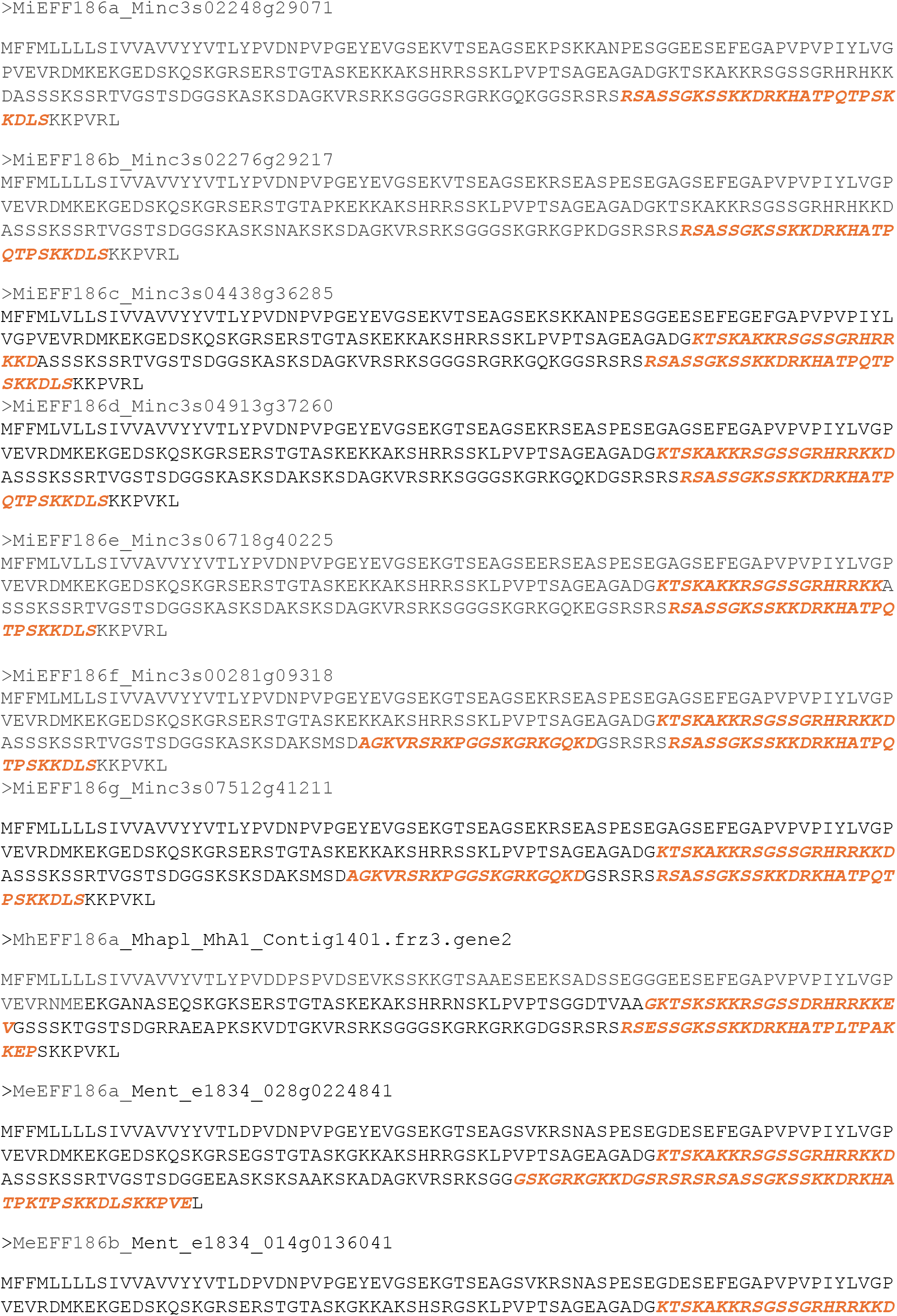
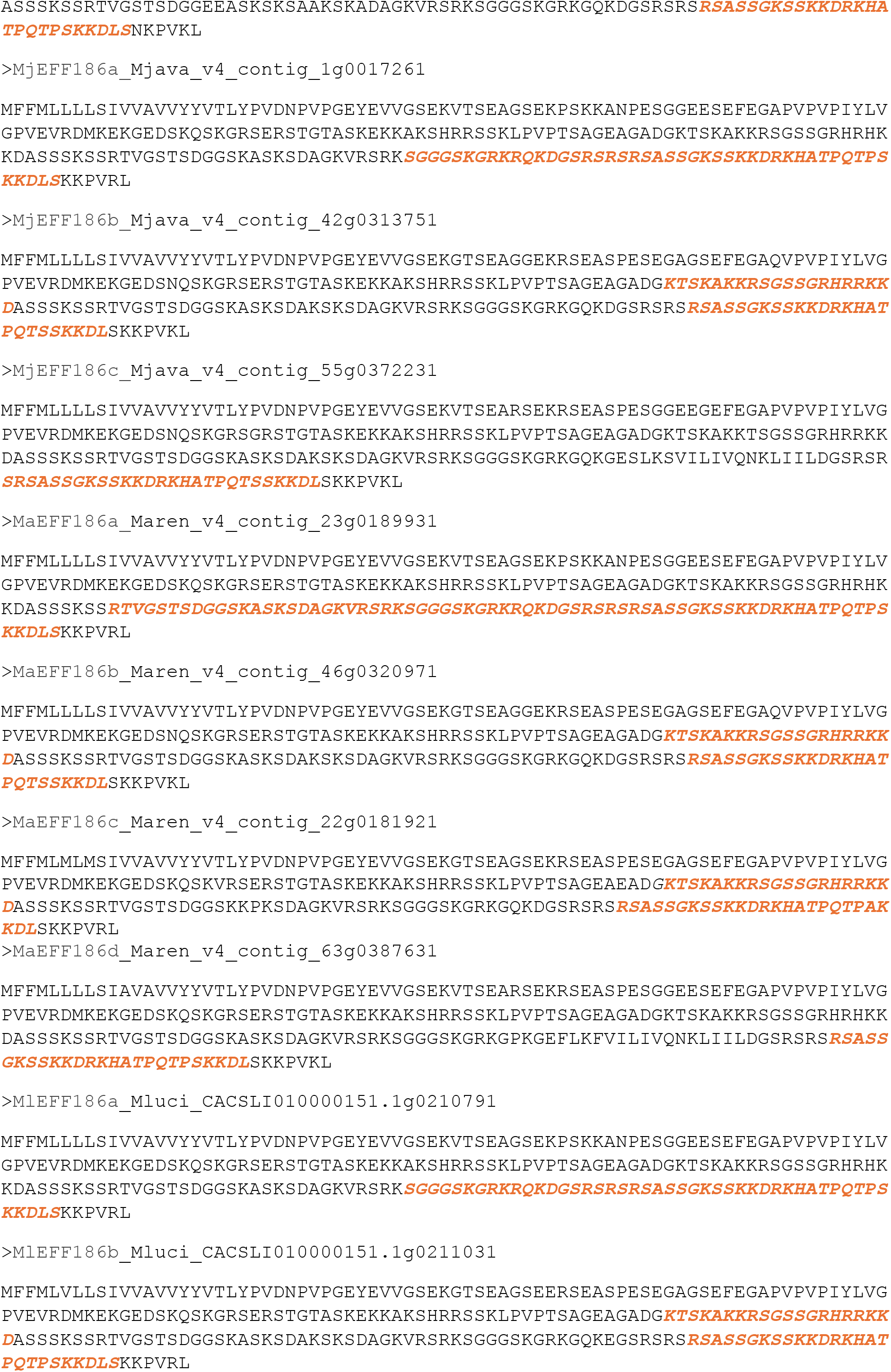
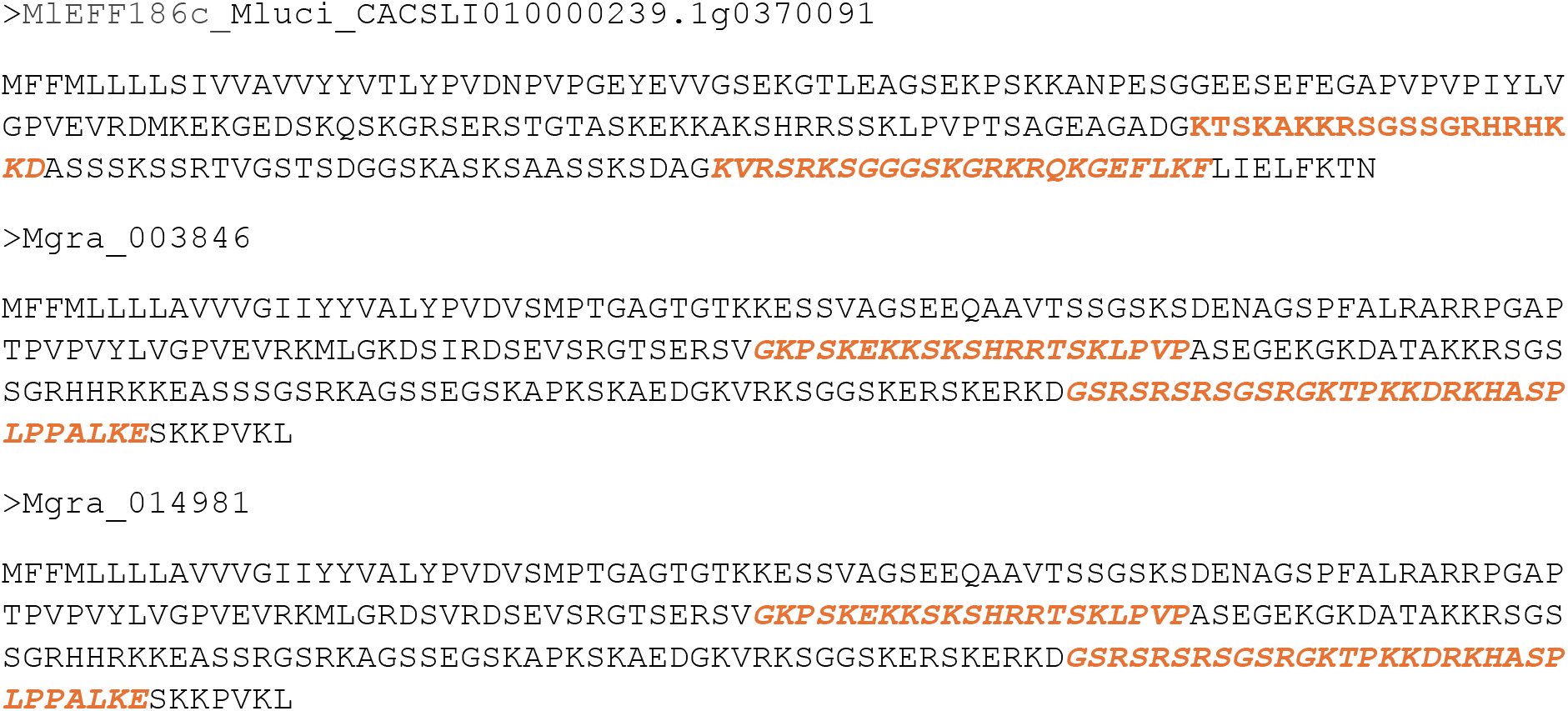
Amino acid sequences of EFF186 effector proteins identified in RKN species. Nucleolar Localisation Signal (NoLS) sequences are indicated in italics and orange.

**Supplementary Fig. S4.**
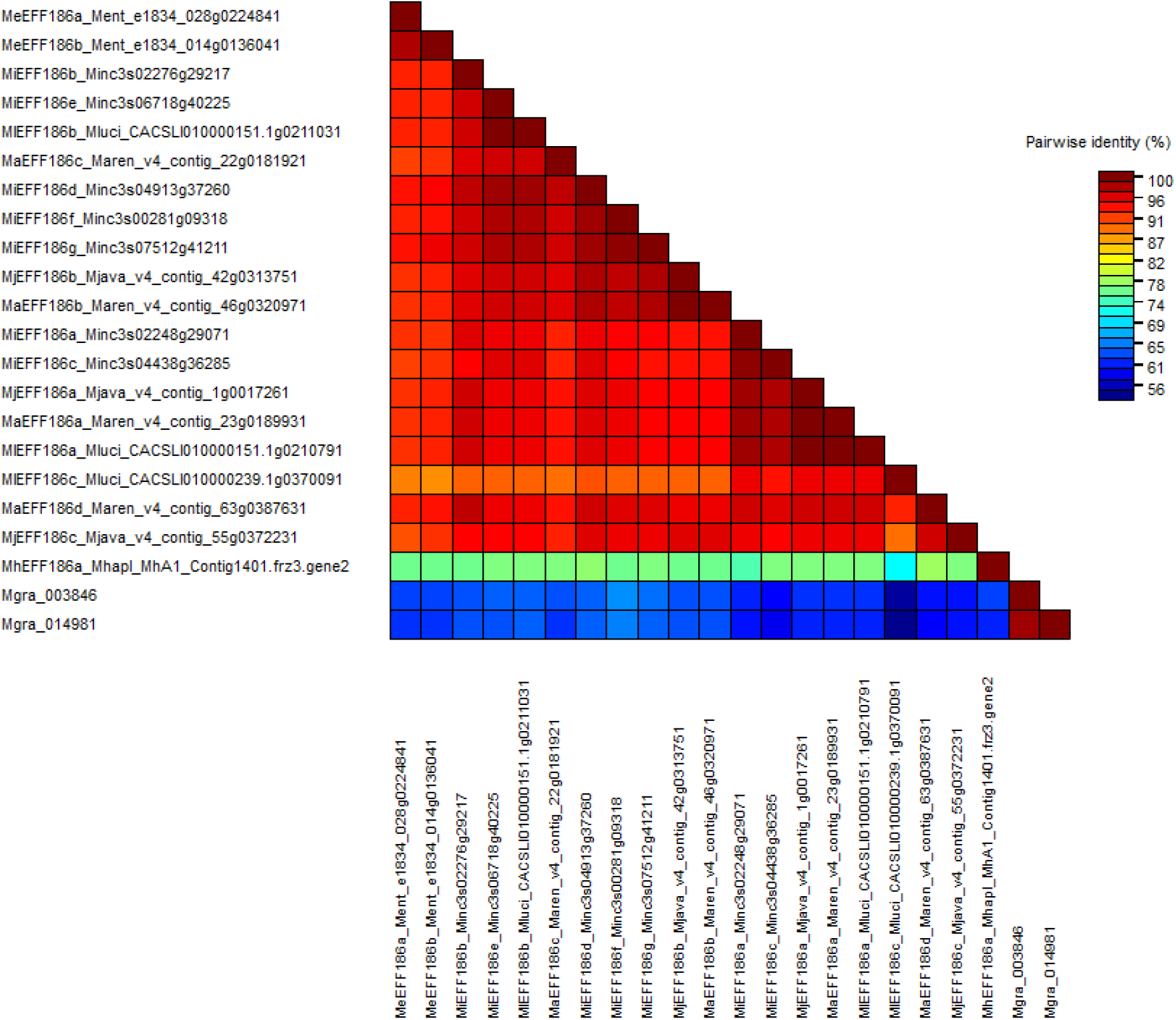
Pairwise sequence identity matrix for root-knot nematodes EFF186 protein sequences.

**Supplementary Fig. S5.**
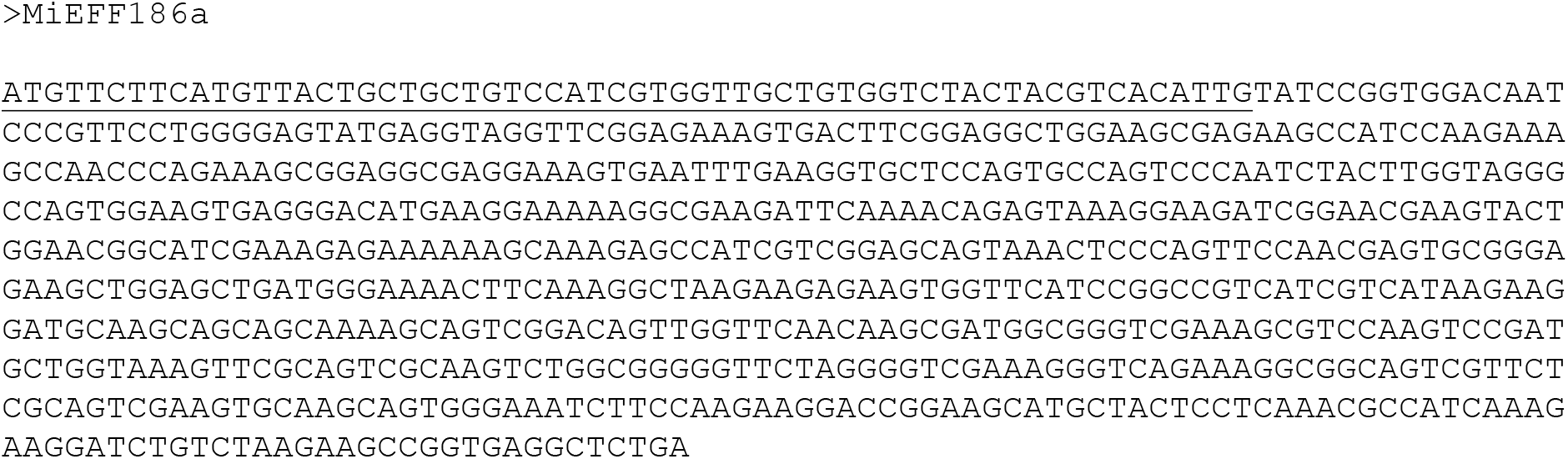
Coding sequences (CDS) of *M. incognita MiEFF186a*. The sequence coding for the signal peptide for secretion is underlined.

**Supplementary Fig. S6.**
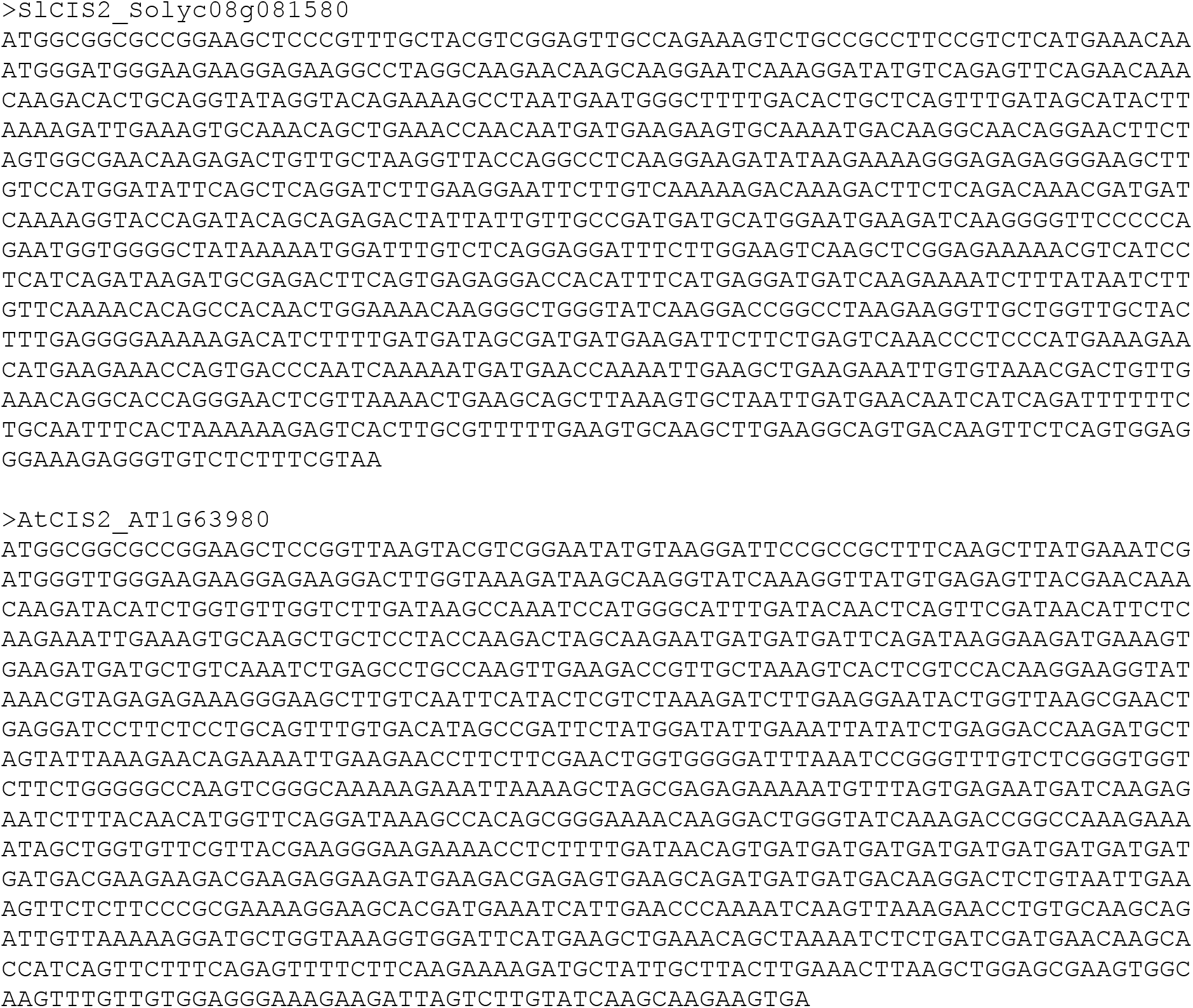
Coding sequences (CDS) of *Solanum lycopersicum* and *Arabidopsis thaliana* CIS2/GDP1.

**Supplementary Fig. S7.**
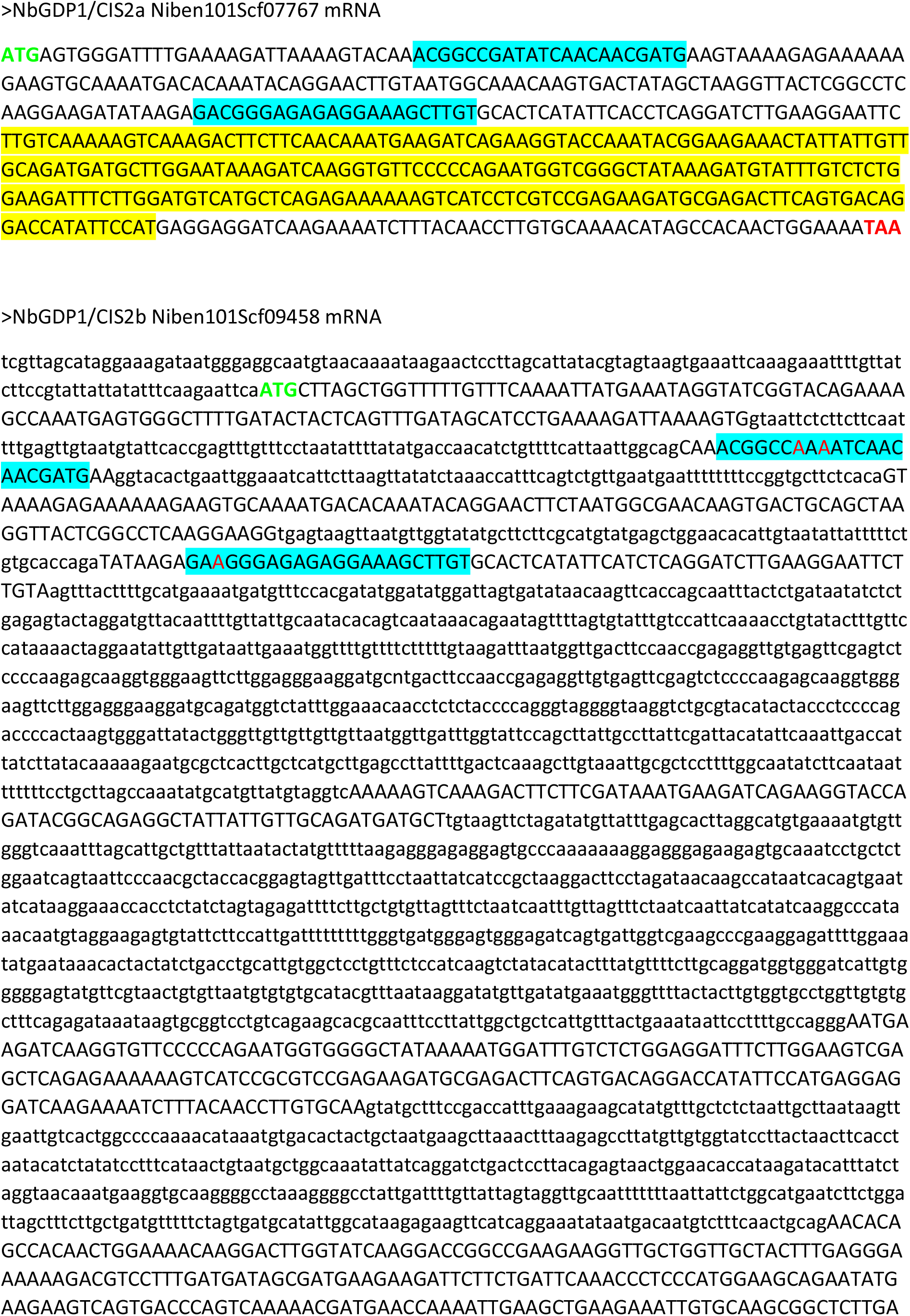
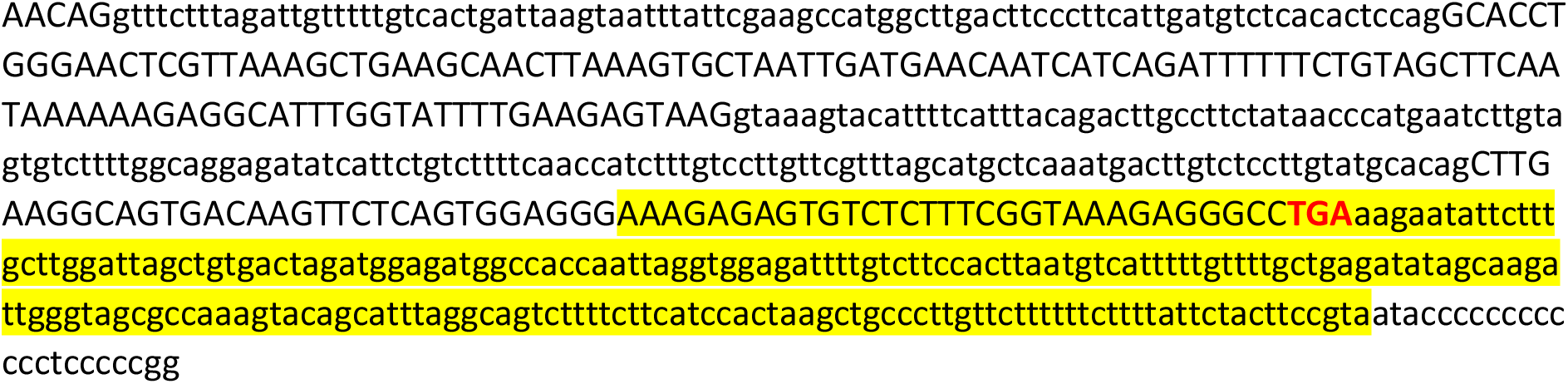
Nucleotide sequences of *Nicotiana benthamiana CIS2/GDP1* genes. The mRNA sequences of GDP1/CIS2 are shown. The open reading frames (ORF) appear in upper case and the untranslated regions (UTR) and introns in lower case. The sequences selected by the VIGS tool to target CIS2/GDP1 are highlighted in yellow. Primers used to validate silencing of GDP1/CIS2 genes by RT-qPCR are highlighted in blue.

**Supplementary Fig. S8.**
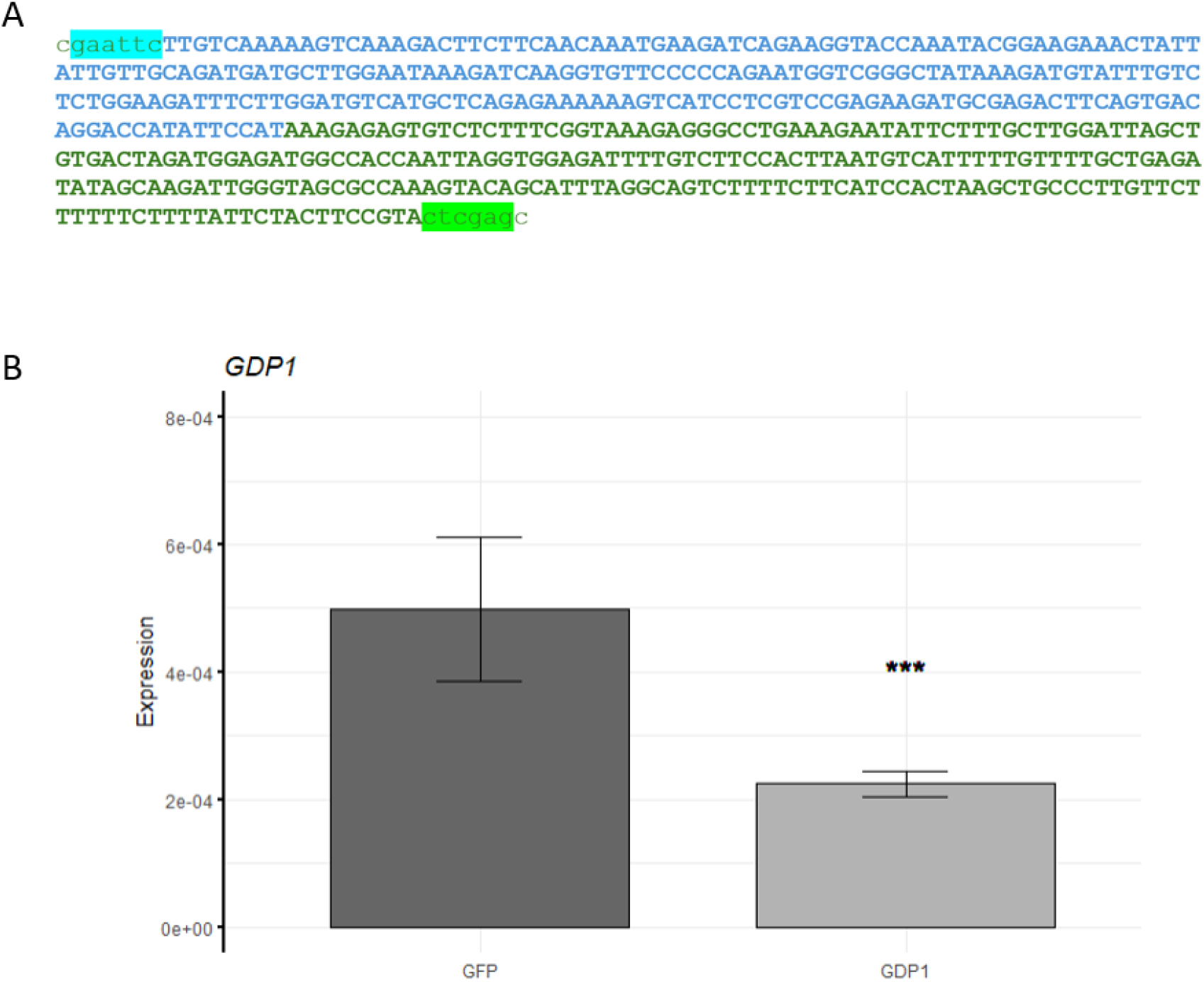
Silencing of the *NbCIS2/GDP1* genes through VIGS. (A) A chimeric sequence was introduced into the TRV2 RNA to silence the two *NbCIS2/GDP1* genes through VIGS. Parts of the sequence in blue and green were selected with the VIGS Tool (https://vigs.solgenomics.net/) for the specific targeting of *NbGDP1/CIS2a* and *NbGDP1/CIS2b*, respectively. The EcoRI and XhoI restriction sites used for cloning into the TRV2 vector are highlighted in blue and green, respectively. (b) RT-qPCR showing *GDP1/CIS2* genes expression in *N. benthamiana* leaves after silencing by VIGS.

**Supplementary Fig. S9.**
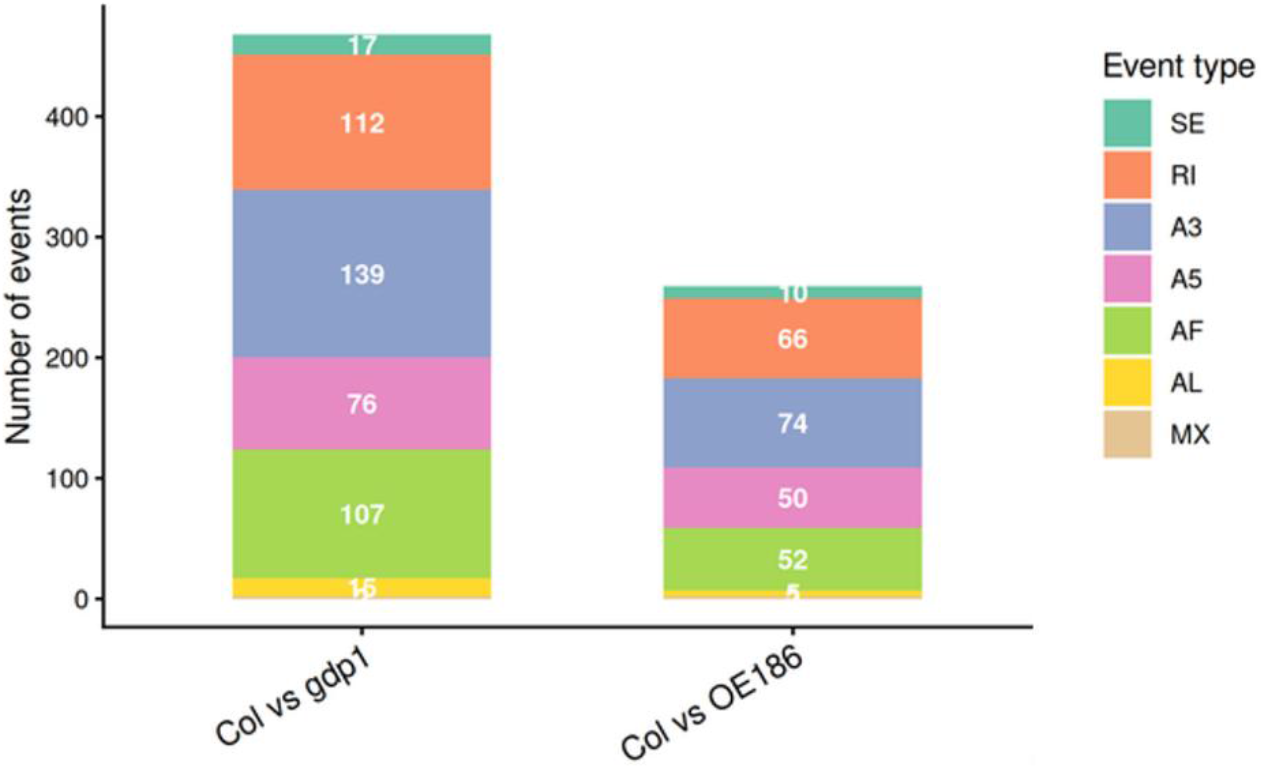
SUPPA2 analysis identified differential alternative splicing events in the *Arabidopsis gdp1.2* mutant and the *MiEFF186a*-overexpressing line.

**Table S1.**
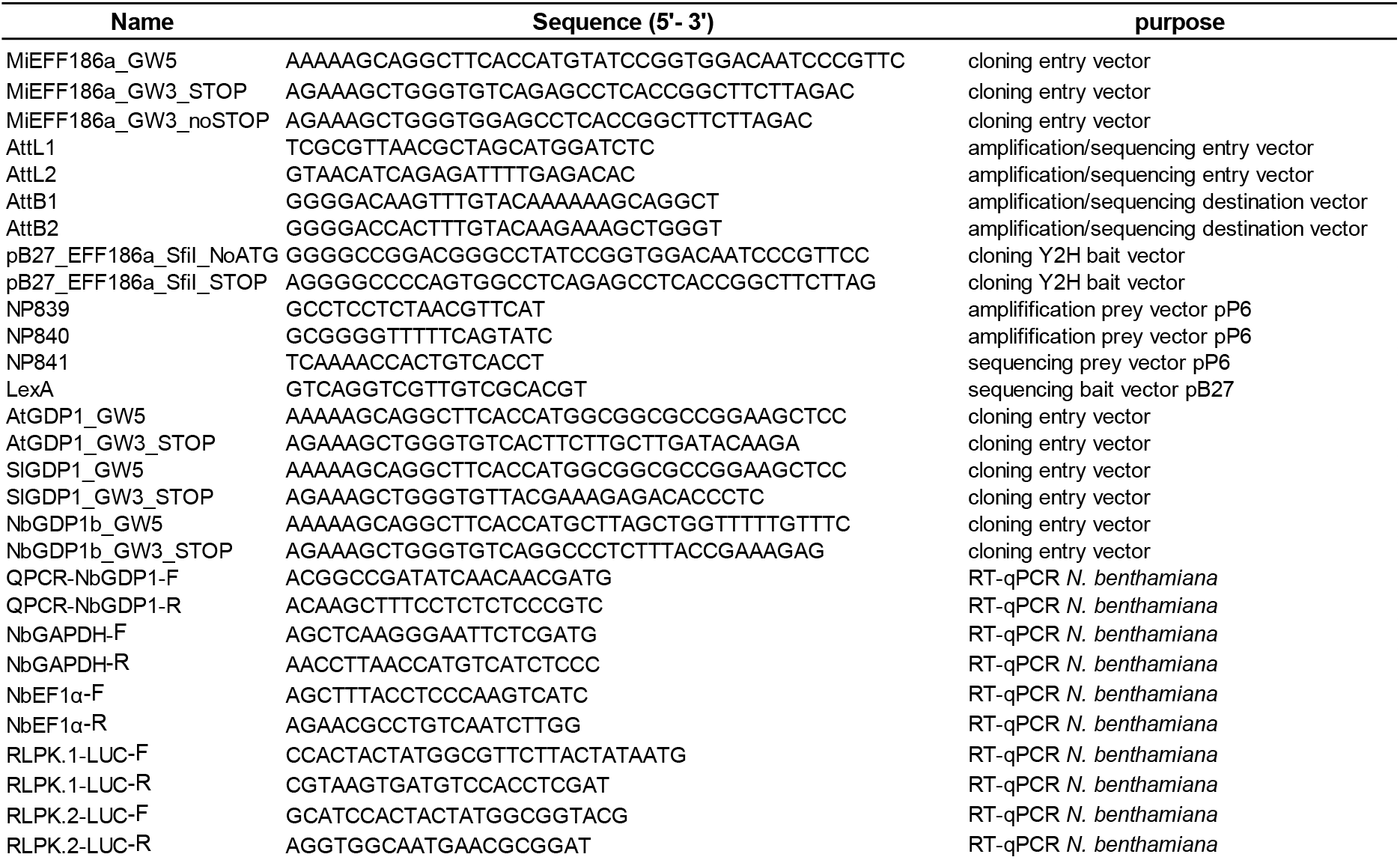
Specific primers used in this study.

## Notes

### Competing Interest Statement

The authors have declared no competing interest.

## REFERENCES

1. Almagro Armenteros, J. J., Tsirigos, K. D., Sønderby, C. K., Petersen, T. N., Winther, O., Brunak, S., von Heijne, G., & Nielsen, H. (2019) SignalP 5.0 improves signal peptide predictions using deep neural networks. Nature Biotechnology, 37(4), 420–423. doi: 10.1038/s41587-019-0036-z

2. Bali, S., & Gleason, C. (2024) Unveiling the Diversity: Plant Parasitic Nematode Effectors and Their Plant Interaction Partners. Molecular Plant-Microbe Interactions, 37(3), 179–189. doi:10.1094/MPMI-09-23-0124-FI

3. Betz, R., Heidt, S., Figueira-Galán, D., Hartmann, M., Langner, T., Requena, N. (2024) Alternative splicing regulation in plants by SP7-like effectors from symbiotic arbuscular mycorrhizal fungi. Nature Communications, 15(1):7107. doi: 10.1038/s41467-024-51512-5

4. Caillaud, M.-C., & Favery, B. (2016). *In vivo* imaging of microtubule organization in dividing giant cell. In Plant Cell Division: Methods and Protocols (M.-C. Caillaud (ed.); pp. 137–144). Springer New York. doi: 10.1007/978-1-4939-3142-2_11

5. Calia, G., Porracciolo, P., Chen, Y., Kozlowski, D., Schuler, H., Cestaro, A., Quentin, M., Favery, B., Danchin, E.G.J. & Bottini, S. (2024) Identification and characterization of specific motifs in effector proteins of plant parasites using MOnSTER. Communications Biology, 7(1):850. doi: 10.1038/s42003-024-06515-9

6. Camborde, L., Kiselev, A., Pel, M. J. C., Le Ru, A., Jauneau, A., Pouzet, C., Dumas, B., & Gaulin, E. (2022). An oomycete effector targets a plant RNA helicase involved in root development and defense. New Phytologist, 233(5), 2232–2248. doi:10.1111/nph.17918

7. Chen, C., Hu, W., Chen, C., Qin, X., Yu, J., Jiang, Y., Hu, J., Zeng, S., Wang, R., Yang, X. & Wang, X. (2025) A nematode effector, MiMSP8, targets U2AF, a subunit of a splicing factor involved in host pre-mRNA splicing, thereby promoting parasitism. Plant Cell and Environment, 48(9):6847–6864. doi: 10.1111/pce.15661

8. Choi, I., Jeon, Y., Yoo, Y., Cho, H. S. & Pai, H. S. (2020) The in vivo functions of ARPF2 and ARRS1 in ribosomal RNA processing and ribosome biogenesis in *Arabidopsis*. Journal of Experimental Botany, 71: 2596–2611. doi: 10.1093/jxb/eraa019

9. Da Rocha, M., Bournaud, C., Dazenière, J., Thorpe, P., Bailly-Bechet, M., Pellegrin, C., Péré, A., Grynberg, P., Perfus-Barbeoch, L., Eves-van den Akker, S. & Danchin, E.G.J. (2021) Genome expression dynamics reveal the parasitism regulatory landscape of the root-knot nematode *Meloidogyne incognita* and a promoter motif associated with effector genes. Genes , 12(5):771. doi: 10.3390/genes12050771

10. Du, K., Peng, D., Wu, J., Zhu, Y., Jiang, T., Wang, P., Chen, X., Jiang, S., Li, X., Cao, Z., Fan, Z. & Zhou, T. (2024) Maize splicing-mediated mRNA surveillance impeded by sugarcane mosaic virus-coded pathogenic protein NIa-Pro. Science Advances, 10(34):eadn3010. doi: 10.1126/sciadv.adn3010

11. Favery, B., Dubreuil, G., Chen, M. S., Giron, D., & Abad, P. (2020) Gall-inducing parasites: convergent and conserved strategies of plant manipulation by insects and nematodes. Annual Review of Phytopathology, 58, 1–22. doi: 10.1146/annurev-phyto-010820-012722

12. Gao, M., Lozano-Durán, R. & Pott, D.M., (2026) New insights into microbial manipulation of the plant spliceosome. Journal of Experimental Botany, erag139. doi: 10.1093/jxb/erag139

13. Godinho, D.P., Yanez, R.J.R., Duque, P. (2025) Pathogen-responsive alternative splicing in plant immunity. Trends in Plant Science, 30(6):615–628, doi: 10.1016/j.tplants.2024.11.010

14. Grynberg, P., Coiti Togawa, R., Dias de Freitas, L., Antonino, J.D., Rancurel, C., Mota do Carmo Costa, M., Grossi-de-Sa, M.F., Miller, R.N., Brasileiro, A.C.M., Messenberg Guimaraes, P. & Danchin, E.G. (2020) Comparative genomics reveals novel target genes towards specific control of plant-parasitic nematodes. Genes 11: 1347. doi:10.3390/genes11111347

15. Hewezi, T. (2024) Phytopathogens reprogram host alternative mRNA splicing. Annual Review of Phytopathology, 173–192. doi: 10.1146/annurev-phyto-121423-041908

16. Huang, J., Gu, L., Zhang, Y., Yan, T., Kong, G., Kong, L., Guo, B., Qiu, M., Wang, Y., Jing, M., Xing, W., Ye, W., Wu, Z., Zhang, Z., Zheng, X., Gijzen, M., Wang, Y., & Dong, S. (2017). An oomycete plant pathogen reprograms host pre-mRNA splicing to subvert immunity. Nature Communications, 8(1). doi: 10.1038/s41467-017-02233-5

17. Huang, J., Lu, X., Wu, H., Xie, Y., Peng, Q., Gu, L., Wu, J., Wang, Y., Reddy, A. S. N., & Dong, S. (2020) *Phytophthora* effectors modulate genome-wide alternative splicing of host mRNAs to reprogram plant immunity. Molecular Plant, 13(10), 1470–1484. doi: 10.1016/j.molp.2020.07.007

18. Jabre, I., Khokhar, W., Reddy, A., Wilson, C. & Syed, N.H. (2026) Alternative splicing in plants: emerging order from chaos. Trends in Plant Science, 3, 1014–1029. doi: 10.1016/j.tplants.2026.04.001

19. Jaouannet, M., Nguyen, C.-N., Quentin, M., Jaubert-Possamai, S., Rosso, M.-N., & Favery, B. (2018) *In situ* hybridization (ISH) in preparasitic and parasitic stages of the plant-parasitic nematode *Meloidogyne* spp. BIO-PROTOCOL, 8(6). doi:10.21769/bioprotoc.2766

20. Jaouannet, M., Perfus-Barbeoch, L., Deleury, E., Magliano, M., Engler, G., Vieira, P., Danchin, E.G.J., Rocha, M.D., Coquillard, P., Abad, P. & Rosso, M.N. (2012) A root-knot nematode-secreted protein is injected into giant cells and targeted to the nuclei. New Phytol. 2012 Jun;194(4):924–931. doi: 10.1111/j.1469-8137.2012.04164.x

21. Karika, L.O., Cipakova, I., Hronska, L. & Cipak, L. (2026) G-patch proteins: important regulators of pre-mRNA splicing and ribosome biogenesis. Frontiers in Cell Developmental Biology, 14:1750689. doi: 10.3389/fcell.2026.1750689

22. Karimi, M., Inzé, D., & Depicker, A. (2002) GATEWAY vectors for Agrobacterium-mediated plant transformation. Trends in Plant Science, 7(5):193–5. doi: 10.1016/s1360-1385(02)02251-3

23. Kojima, K., Tamura, J., Chiba, H., Fukada, K., Tsukaya, H., & Horiguchi, G. (2018) Two nucleolar proteins, GDP1 and OLI2, function as ribosome biogenesis factors and are preferentially involved in promotion of leaf cell proliferation without strongly affecting leaf adaxial–abaxial patterning in *Arabidopsis thaliana*. Frontiers in Plant Science, 8:, 1–15. doi:10.3389/fpls.2017.02240

24. Kumar, S., Stecher, G., & Tamura, K. (2016) MEGA7: molecular evolutionary genetics analysis version 7.0 for bigger datasets. Molecular Biology and Evolution, 33(7), 1870–1874. doi:10.1093/MOLBEV/MSW054

25. Laloum, T., Martín, G., & Duque, P. (2018) Alternative splicing control of abiotic stress responses. Trends in Plant Science, 23(2), 140–150. doi:10.1016/j.tplants.2017.09.019

26. Lee, S., Seo, Y.E., Choi, J., Yan, X., Kim, T., Choi, D. & Lee, J.H. (2024) Nucleolar actions in plant development and stress responses. Plant Cell and Environment, 47(12):5189–5204. doi: 10.1111/pce.15099

27. Li, Y. & Kou, S. (2025) A *Ralstonia solanacearum* effector targets splicing factor SR34a to reprogram alternative splicing and regulate plant immunity. Plants, 14, 534. doi: 10.3390/Plants14040534

28. Lu, X., Yang, Z., Song, W., Miao, J., Zhao, H., Ji, P., Li, T., Si, J., Yin, Z., Jing, M., Shen, D., & Dou, D. (2023). The Phytophthora sojae effector PsFYVE1 modulates immunity-related gene expression by targeting host RZ-1A protein. Plant Physiology, 191(2), 925–945. doi:10.1093/plphys/kiac552

29. Maekawa, S., Ueda, Y. & Yanagisawa, S. (2018) Overexpression of a Brix domain-containing ribosome biogenesis factor ARPF2 and its interactor ARRS1 causes morphological changes and lifespan extension in *Arabidopsis thaliana*. Frontiers in Plant Science, 9: 1177. doi: 10.3389/fpls.2018.01177

30. Mejias, J., Bazin, J., Truong, N. M., Chen, Y., Marteu, N., Bouteiller, N., Sawa, S., Crespi, M. D., Vaucheret, H., Abad, P., Favery, B., & Quentin, M. (2021) The root-knot nematode effector MiEFF18 interacts with the plant core spliceosomal protein SmD1 required for giant cell formation. New Phytologist, 229(6), 3408–3423. doi: 10.1111/nph.17089

31. Mejias, J., Chen, Y., Bazin, J., Truong, N. M., Mulet, K., Noureddine, Y., Jaubert-Possamai, S., Ranty-Roby, S., Soulé, S., Abad, P., Crespi, M. D., Favery, B., & Quentin, M. (2022) Silencing the conserved small nuclear ribonucleoprotein *SmD1* target gene alters susceptibility to root-knot nematodes in plants. Plant Physiology, 189(3), 1741–1756. doi: 10.1093/plphys/kiac155

32. Mitchum, M., Rocha, R., Huang, G. Maier, T., Baum, T. & Hussey, R. (2023) Genome-guided reanalysis of root-knot nematode *Meloidogyne incognita* esophageal gland cell-enriched sequence tag libraries: a resource for the discovery of novel effectors. PhytoFrontiers, 3: 435–441. doi: 10.1094/PHYTOFR-09-22-0099-A

33. Nakai, K., & Horton, P. (1999). PSORT: A program for detecting sorting signals in proteins and predicting their subcellular localization. Trends in Biochemical Sciences, 24(1), 34–35. doi: 10.1016/S0968-0004(98)01336-X

34. Nicaise, V., Joe, A., Jeong, B. R., Korneli, C., Boutrot, F., Westedt, I., Staiger, D., Alfano, J. R., & Zipfel, C. (2013) *Pseudomonas* HopU1 modulates plant immune receptor levels by blocking the interaction of their mRNAs with GRP7. EMBO Journal, 32(5), 701–712. doi: 10.1038/emboj.2013.15

35. Ozdemir, S., Piya, S., Lopes-Caitar, V.S., Coffey, N., Rice, J.H. & Hewezi, T. (2024) Local and systemic transcriptome and spliceome reprogramming induced by the root-knot nematode *Meloidogyne incognita* in tomato. Horticulture Research, 11, uhae206, doi: 10.1093/hr/uhae206

36. Plaschka, C., PC, L., Charenton, C., & Nagai, K. (2018) Prespliceosome structure provides insights into spliceosome assembly and regulation. Nature, 559(7714), 419–422. doi: 10.1038/s41586-018-0323-8

37. Reddy, A. S. N. (2007) Alternative splicing of pre-messenger RNAs in plants in the genomic era. Annual Review of Plant Biology, 58, 267–294. doi: 10.1146/annurev.arplant.58.032806.103754

38. Rigo, R., Bazin, J., Crespi, M., & Charon, C. (2019) Alternative splicing in the regulation of plant-microbe interactions. Plant and Cell Physiology, 60(9), 1906–1916. doi:10.1093/pcp/pcz086

39. Rocha RO, Hussey RS, Pepi LE, Azadi P, Mitchum MG. (2023) Discovery of novel effector protein candidates produced in the dorsal gland of adult female root-knot nematodes. Molecular Plant Microbe Interaction, 36(6):372–380. doi: 10.1094/MPMI-11-22-0232-R

40. Rutter, W. B., Franco, J., & Gleason, C. (2022) Rooting out the mechanisms of root-knot nematode– plant interactions. Annual Review of Phytopathology, 1–34. doi: 10.1146/annurev-phyto-021621-120943

41. Rutter, W.B., Hewezi, T., Abubucker, S., Maier, T.R., Huang, G., Mitreva, M., Hussey, R.S. & Baum, T.J. (2014) Mining novel effector proteins from the esophageal gland cells of Meloidogyne incognita. Molecular Plant-Microbe Interactions, 27: 965–974. doi: 10.1094/MPMI-03-14-0076-R

42. Scott, M. S., Troshin, P. V., & Barton, G. J. (2011) NoD: A Nucleolar localization sequence detector for eukaryotic and viral proteins. BMC Bioinformatics, 12. doi: 10.1186/1471-2105-12-317

43. Soulé, S., Huang, K., Mulet, K., Mejias, J., Bazin, J., Truong, N.M., Kika, J.L., Jaubert, S., Abad, P., Zhao, J., Favery, B. & Quentin, M. (2024) The root-knot nematode effector MiEFF12 targets the host ER quality control system to suppress immune responses and allow parasitism. Molecular Plant Pathology, 25(7):e13491. doi: 10.1111/mpp.13491

44. Tognacca, R.S., Rodríguez, F.S., Aballay, F.E., Cartagena, C.M., Servi, L., Petrillo, E. (2023) Alternative splicing in plants: current knowledge and future directions for assessing the biological relevance of splice variants. Journal of Experimental Botany, 74(7):2251–2272. doi: 10.1093/jxb/erac431

45. Trivedi, R., & Nagarajaram, H. A. (2022). Intrinsically disordered proteins: an overview. International Journal of Molecular Sciences, 23(22), 1–30. doi: 10.3390/ijms232214050

46. Verma, A., Lee, C., Morriss, S., Odu, F., Kenning, C., Rizzo, N., Spollen, W. G., Lin, M., McRae, A. G., Givan, S. A., Hewezi, T., Hussey, R., Davis, E. L., Baum, T. J., & Mitchum, M. G. (2018). The novel cyst nematode effector protein 30D08 targets host nuclear functions to alter gene expression in feeding sites. New Phytologist, 219(2), 697–713. doi: 10.1111/nph.15179

47. Vijayapalani, P., Hewezi, T., Pontvianne, F. & Baum, T.J. (2018) An effector from the cyst nematode *Heterodera schachtii* derepresses host rRNA genes by altering histone acetylation. The Plant Cell, 30(11):2795–2812. doi: 10.1105/tpc.18.00570

48. Wang, L., Chen, M., Zhu, F., Fan, T., Zhang, J., & Lo, C. (2020) Alternative splicing is a *Sorghum bicolor* defense response to fungal infection. Planta, 251(1), 1–13. doi:10.1007/s00425-019-03309-w

49. Yu, H., Shi, X., Ning, N., Wu, H., Mei, J., Gu, X., Ruan, H., Zhang, M., Li, Z., Ma, S. & Liu, W. (2025) The *Exserohilum turcicum* effector EtEC81 reprograms alternative splicing in maize and activates immunity. Cell Reports, 44(4):115501. doi: 10.1016/j.celrep.2025.115501

50. Zhao, Z., Dent, C., Liang, H., Lv, J., Shang, G., Liu, Y., Feng, F., Wang, F., Pang, J., Li, X., Ma, L., Li, B., Sureshkumar, S., Wang, J. W., Balasubramanian, S., & Liu, H. (2022) CRY2 interacts with CIS1 to regulate thermosensory flowering via *FLM* alternative splicing. Nature Communications, 13(1), 7045. doi: 10.1038/s41467-022-34886-2

51. Zhao, J., Mejias, J., Quentin, M., Chen, Y., de Almeida-Engler, J., Mao, Z., Sun, Q., Liu, Q., Xie, B., Abad, P., Favery, B. & Jian, H. (2020) The root-knot nematode effector MiPDI1 targets a stress-associated protein (SAP) to establish disease in Solanaceae and Arabidopsis. New Phytologist, 228(4):1417–1430. doi: 10.1111/nph.16745

